# Nascent Chain Ubiquitination is Uncoupled from Degradation to Enable Protein Maturation

**DOI:** 10.1101/2023.10.09.561585

**Authors:** Xia Li, Malaiyalam Mariappan

## Abstract

A significant proportion of nascent proteins undergo polyubiquitination on ribosomes in mammalian cells, yet the fate of these proteins remains elusive. The ribosome-associated quality control (RQC) is a mechanism that mediates the ubiquitination of nascent chains on stalled ribosomes. Here, we find that nascent proteins ubiquitinated on stalled ribosomes by the RQC E3 ligase LTN1 are insufficient for proteasomal degradation. Our biochemical reconstitution studies reveal that ubiquitinated nascent chains are promptly deubiquitinated in the cytosol upon release from stalled ribosomes, as they are no longer associated with LTN1 E3 ligase for continuous ubiquitination to compete with cytosolic deubiquitinases. These deubiquitinated nascent chains can mature into stable proteins. However, if they misfold and expose a degradation signal, the cytosolic quality control recognizes them for re-ubiquitination and subsequent proteasomal degradation. Thus, our findings suggest that cycles of ubiquitination and deubiquitination spare foldable nascent proteins while ensuring the degradation of terminally misfolded proteins.

## INTRODUCTION

Cells face fundamental challenges in accurately selecting aberrant proteins for degradation while sparing newly synthesized proteins since both often exhibit similar biophysical characteristics such as exposed hydrophobic sequences^1^. Inefficient recognition and removal of misfolded proteins can lead to the buildup of toxic species, contributing to various neurodegenerative diseases^2,3^. Conversely, prematurely flagging newly synthesized proteins based solely on their initial biophysical properties can lead to wasted cellular energy and disruption of proteostasis. This issue is particularly pronounced during protein synthesis, as a significant portion of nascent proteins on ribosomes undergo polyubiquitination, even under unstressed conditions in both yeast and mammalian cells^4–6^. It has been debated whether all ubiquitinated nascent proteins are degraded because the constant flux of ubiquitinated proteins could substantially burden the proteasome capacity and delay the degradation of toxic misfolded proteins^7–10^. Additionally, the indiscriminate degradation of nascent chains ubiquitinated on ribosomes could lead to wastage of cellular energy, given that each peptide bond formation consumes an energy equivalent of approximately ∼5 ATP molecules^10,11^. Hence, gaining insight into the cellular mechanisms that distinguish terminally misfolded proteins for degradation while allowing folding-competent nascent proteins to mature holds promise in identifying therapeutic strategies for addressing human diseases linked to proteostasis breakdown.

The Ribosome-Associated Quality Control (RQC) is a mechanism responsible for ubiquitinating nascent proteins when ribosomes encounter stalls during translational elongation^12–16^. These stalls can occur under various pathological conditions, such as a deficiency in aminoacyl tRNAs or due to aberrant mRNAs^17^. Furthermore, studies have demonstrated that ribosome stalling can happen under physiological conditions. For instance, specific amino acid sequences, like arrest peptides (APs) and transmembrane domains^18–24^, interact with the components in the ribosome exit tunnel components, leading to the induction of ribosome stalling. These physiological stalls play a critical role in regulating processes such as protein expression, targeting, and folding^25–28^. According to the current model^13–15,29–31^, RQC factors detect stalled ribosomes and facilitate their separation into 60S and 40S subunits. The E3 ligase LTN1 (Listerin)^12^ is responsible for ubiquitinating nascent chains associated with the 60S subunits^32–34^, a process facilitated by the protein NEMF^16,35–38^. LTN1 can also access and mediate ubiquitination ribosome-stalled nascent chains at the Sec61 translocon of the endoplasmic reticulum (ER) ^39–42^. Unlike other protein quality control systems that typically recognize substrates, LTN1 recognizes 60S ribosome subunits harboring peptidyl tRNAs bound to NEMF and mediates the ubiquitination of nascent chains regardless of their folding status^43^. The p97 ATPase is proposed to facilitate the extraction of ubiquitinated nascent chains from 60S subunits, subsequently directing them toward proteasomal degradation^16,37,44^.

Despite significant progress in our understanding of the structural and mechanistic aspects of stalled ribosome recognition and nascent chain ubiquitination, the ultimate and critical stage of protein degradation remains poorly elucidated. Crucially, we understand little about the specific features that dictate the proteasomal degradation of ubiquitinated nascent chains arising from stalled ribosomes. Pioneering studies employing model substrates with aberrant mRNAs, which induce ribosome stalling, played a pivotal role in the identification and characterization of RQC factors^13–15,29,30,45^. However, these mRNAs often lead to the production of aberrant proteins, which prompts questions about the mechanisms of nascent chain degradation. Consequently, it remains unclear whether these faulty proteins are also targeted by cytosolic quality control for ubiquitination and subsequent proteasomal degradation.

To address these fundamental questions, we investigated the endogenous stalling substrate XBP1u, a pivotal player in the unfolded protein response (UPR)^46,47^. This substrate encompasses a small hydrophobic region 1 (HR1), a long HR2, and a translational arrest peptide (AP) at the C-terminus^26,48^. XBP1u AP makes extensive contact with the ribosomal exit tunnel components and forms a unique turn in proximity to the peptidyl transferase center, ultimately leading to translational arrest^49^. We and others have previously shown that the translationally arrested XBP1u mRNA ribosome-nascent chains (RNCs) expose a hydrophobic region 2 (HR2) for the recognition by the signal recognition particle (SRP) and targeting to the Sec61 translocon at the ER membrane ^50,51^. In our investigation of XBP1u, we discovered that ubiquitinated nascent chains, catalyzed by LTN1 on stalled ribosomes, undergo rapid deubiquitination by cytosolic deubiquitinases upon their release. Their proteasomal degradation relies entirely on whether deubiquitinated nascent chains expose a degradation signal in the cytosol for subsequent re-ubiquitination. Furthermore, we establish that while LTN1-mediated ubiquitination on ribosomes is dispensable for nascent chain degradation, it plays a crucial role in liberating nascent chains from stalled ribosomes. Overall, our results suggest that the separation of ubiquitination of nascent chains on ribosomes from immediate proteasomal degradation offers a chance for the maturation of folding-competent proteins while also preventing premature degradation, thus conserving energy expended on protein synthesis.

## RESULTS

### LTN1 and NEMF are recruited to stalled ribosomes housing an endogenous staller protein

To determine the fate of ubiquitinated nascent chains on ribosomes, we investigated the degradation of the endogenous ribosome staller protein XBP1u. It induces ribosomal stalling during translation elongation through its C-terminus arrest peptide (AP)^26^ (Figure 1A). To identify protein quality control factors recruited to stalled ribosome-bound nascent chains (RNCs) of XBP1u, we performed an unbiased pull-down of FLAG-tagged XBP1u from 293T cells and identified its associated proteins through mass spectrometry. Figure 1B demonstrates a selective enrichment of ribosomal proteins in XBP1u immunoprecipitation (IPs) compared to IP from control non-transfected cells, supporting the association of XBP1u with stalled ribosomes in cells (Table 1). Consistent with our previous findings^50^, we detected peptides derived from Sec61 translocon components from XBP1u IP, but they were nearly absent in the control IP (Figure 1B and Table 1). Importantly, peptides generated from RQC factors of NEMF and LTN1 E3 ligase were exclusively found in XBP1u IPs.

**Figure 1.**
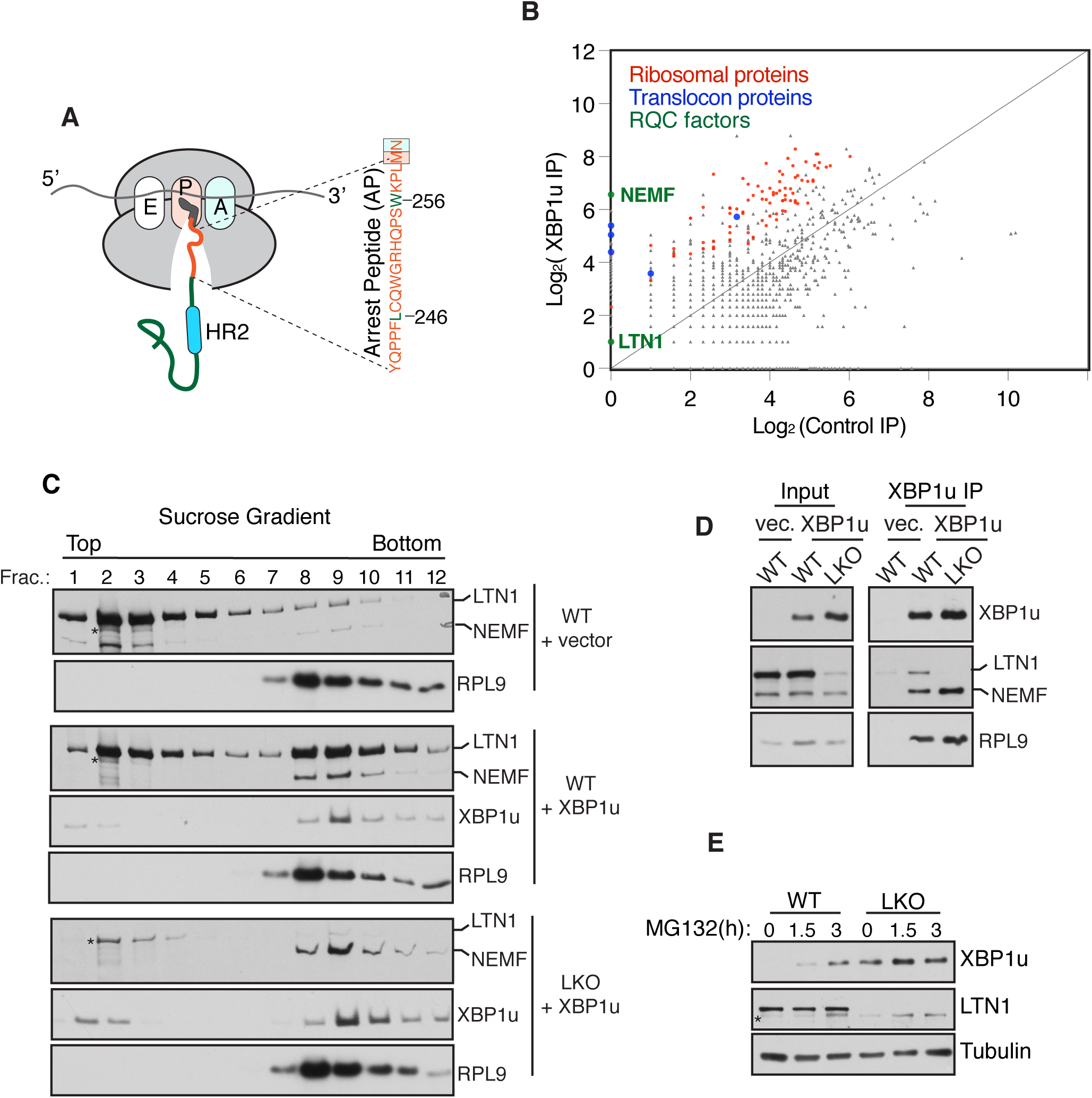
LTN1 and NEMF are recruited to arrest peptide-induced stalled ribosomes in cells. (A) A schematic diagram illustrates the ribosome stalling induced by XBP1u arrest peptide (AP). The incorporation of the last asparagine (N) residue is inhibited due to the stall. The mutation of L246 and W256 to alanine residues relieves the stall. HR2 denotes hydrophobic region 2. (B) Scatter plot comparing the total mass spectrum between control immunoprecipitation (IP) and XBP1u IP. The plot displays the data on a log2 scale representing the total mass spectrum for each protein. (C) WT 293T cells or LTN1 knock-out (LKO) cells expressing empty vector or XBP1u were fractionated by 10-30% sucrose density gradient, and the fractions were analyzed by immunoblotting for the indicated proteins. Star symbol indicates a non-specific band detected by LTN1 antibody. (D) The indicated cells expressing empty vector or FLAG-tagged XBP1u were lysed and immunoprecipitated for XBP1u using anti-FLAG beads and analyzed by immunoblotting. (E) WT and LKO cells were treated with MG132 (20μM) for the indicated time points and analyzed by immunoblotting. See also **Figure S1.**

We validated whether stalled RNCs of XBP1u recruit NEMF and LTN1 by sucrose density gradient fractionation. Immunoblotting of sucrose gradient fractions from non-transfected wild type (WT) 293T cells revealed that both LTN1 and NEMF primarily co-fractionated in the non-ribosomal top fractions, with only a minor fraction localized to the ribosomal fractions (8 to 10) (Figure 1C). In cells expressing XBP1u, as expected, translationally stalled XBP1u co-migrated with ribosomes. Additionally, a notable portion of LTN1, along with nearly the entire population of NEMF, shifted to ribosomal fractions, indicating that XBP1u-induced ribosome stalling initiates the recruitment of these factors. The binding of NEMF to ribosomes appeared independent of LTN1, as its entire population was recruited to stalled XBP1u ribosomes in LTN1 knock-out (LKO) cells generated by CRISPR/Cas9 (Figure 1C). Immunoprecipitation experiments further confirmed the association of XBP1u RNCs with LTN1 and NEMF (Figure 1D).

To investigate if LTN1 E3 ligase activity is required for proteasomal degradation of XBP1u, we probed the stability of endogenous XBP1u in both WT and LKO cells. The endogenous XBP1u was not readily detectable in WT cells (Figure 1E), but inhibition of the proteasome with MG132 led to an accumulation of XBP1u protein, indicating that XBP1u is subjected to proteasomal degradation. Remarkably, levels of XBP1u protein were significantly elevated in LTN1-depleted cells, even in the absence of MG132 treatment (Figure 1E), which is consistent with an earlier study showing LTN1 E3 ligase activity is required for the degradation of XBP1u^21^. Transient expression of recombinant XBP1u exhibited similar behavior to its endogenous counterpart, accumulating in LKO or LTN1-depleted cells using siRNA when compared to control cells (Figure S1A, S1B). This accumulation of XBP1u was restorable through the complementation of WT LTN1, but not with LTN1 lacking the catalytic RING domain (LTN1ΔRING), suggesting that LTN1-mediated ubiquitination plays a crucial role in regulating XBP1u protein levels in cells (Figure S1C). Together, these data suggest that RQC factors are recruited to stalled ribosomes induced by XBP1u AP and appear to mediate the degradation of XBP1u in cells.

### LTN1-mediated ubiquitination is insufficient for the degradation of nascent chains from stalled ribosomes

To first determine whether LTN1 mediates XBP1u ubiquitination on stalled ribosomes, we co-expressed XBP1u along with HA-ubiquitin in both WT and LKO cells. Subsequently, XBP1u was immunoprecipitated using anti-FLAG antibodies, then probing for ubiquitin-conjugated XBP1u via immunoblotting. In WT cells, the ubiquitinated species, characterized by ladder and smear bands, were readily detectable, while their presence in LKO cells was nearly absent (Figure 2A and 2B). Similarly, an arrest peptide (AP) mutant of XBP1u (XBP1u-AP*) that cannot induce ribosome stalling^26^ exhibited ubiquitination but independently of LTN1 activity (Figure 2A and 2C). When probed with a K48-linked ubiquitin chain-specific antibody, it became evident that the polyubiquitinated species of both XBP1u and XBP1u-AP* were enriched in K48-linked ubiquitin chains, known to be optimal for directing proteins for proteasomal degradation (Figure S2A). Pulse-chase and cycloheximide (CHX) chase experiments illustrated that XBP1u underwent rapid turnover in WT cells, while degradation was impeded in LKO cells (Figure 2D and S2B), corroborating the ubiquitination results. In contrast, the ribosomal arrest mutant XBP1u-AP* was subject to degradation in an LTN1-independent manner, as it exhibited a similar turnover rate in both WT and LKO cells (Figure 2E and S2C). Together, these data suggest that XBP1u represents a unique substrate, being ubiquitinated and subsequently degraded irrespective of ribosome stalling.

**Figure 2.**
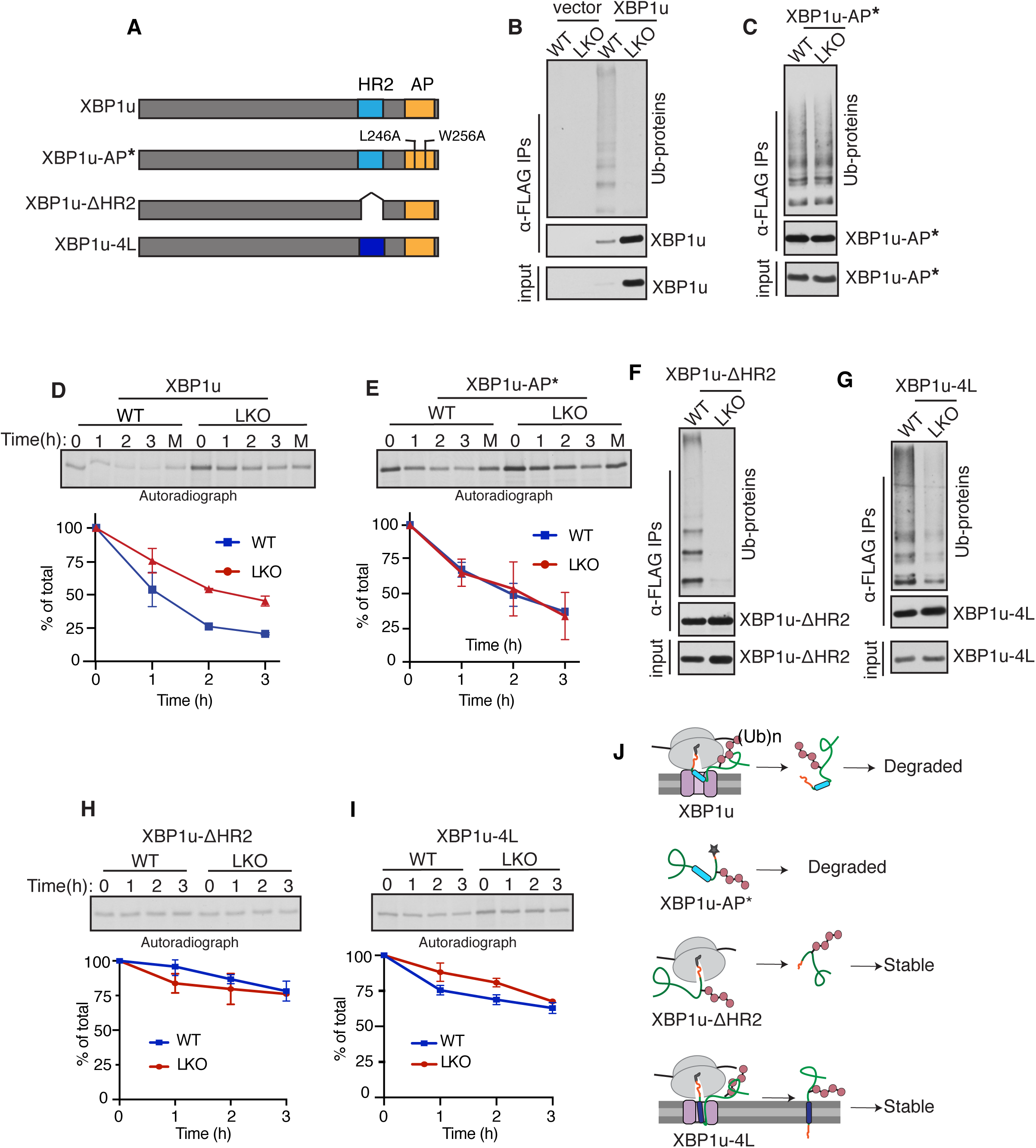
Nascent chain ubiquitination by LTN1 is insufficient for proteasomal degradation. (A) Cartoon illustrating WT XBP1u, the arrest peptide mutant XBP1u-AP*, XBP1u lacking hydrophobic region 2 (HR2), and XBP1u-4L with four hydrophobic leucine residues replacing four less hydrophobic residues in HR2. (B) Empty vector or FLAG-XBP1u is transfected along with HA-ubiquitin into WT or LKO cells and was immunoprecipitated with anti-FLAG beads. The resulting samples were probed for ubiquitinated proteins (Ub-proteins) using an HA antibody and for XBP1u with an anti-FLAG antibody. (C) The indicated cells expressing FLAG-XBP1u-AP* were analyzed as in (B). (D, E) Pulse-chase analysis of WT and LKO cells expressing XBP1u or XBP1u-AP*. The data were quantified from two independent experiments and are represented as means ±SEMs. (F, G) WT or LKO cells expressing XBP1u-ΔHR2 or XBP1u-4L were analyzed as in (B) (H, I) WT or LKO cells expressing XBP1u-ΔHR2 or XBP1u-4L were analyzed by pulse-chase. The data were quantified from two independent experiments and are represented as means ±SEMs. (J) Schematic diagram depicting the critical role of exposed hydrophobicity in XBPu in mediating proteasomal degradation. See also **Figure S2.**

The ubiquitination and subsequent degradation of XBP1u-AP* indicate the presence of a degron mediating proteasomal degradation within XBP1u. Given that XBP1u possesses a long hydrophobic sequence (HR2) and is exposed to the cytosol due to its inability to insert into the ER membrane^50^, we hypothesized that the exposed HR2 may play a primary role in mediating the degradation of XBP1u. To test this, we first removed exposed hydrophobicity in XBP1u by deleting HR2 (XBP1u-ΔHR2) (Figure 2A). Second, we generated the XBP1u-4L construct, in which less hydrophobic amino acids in HR2 were replaced with hydrophobic leucines (Figure 2A). XBP1u-4L is known to efficiently insert into the ER membrane, thus preventing exposure to the cytosol^52^. XBP1u-ΔHR2 and XBP1u-4L were still ubiquitinated in an LTN1-dependent manner, as both contain intact AP at their C-terminus (Figure 2A, 2F, and 2G). Similar LTN1-dependent ubiquitination was observed when HR2 was replaced with a hydrophilic sequence, thus excluding potential effects caused by HR2 deletion (Figure S2D). Despite being ubiquitinated by LTN1, both XBP1u-ΔHR2 and XBP1u-4L remained stable in both WT and LKO cells, as confirmed by pulse-chase and CHX chase assays (Figure 2H, 2I and S2E). Collectively, these data suggest that exposed hydrophobicity plays a critical role in the degradation of nascent chains but not the AP-induced and LTN1-medaited ubiquitination that initially occurs on stalled ribosomes (Figure 2J).

### Nascent chains ubiquitinated by LTN1 are deubiquitinated in the cytosol

The above data suggest a model where XBP1u undergoes two rounds of ubiquitination: first by LTN1 on stalled ribosomes and second through HR2-dependent ubiquitination in the cytosol. To test this, we separated ribosomes from the cytosol fraction using a sucrose cushion and probed for ubiquitin-conjugated XBP1u (Figure 3A). Since the nascent chain remains attached to ribosomes, it would co-sediment with them, whereas released nascent chains would be found in the cytosol. This allowed us to distinguish AP-induced nascent chain ubiquitination on ribosomes from HR2-dependent ubiquitination in the cytosol. Consistent with our model, ubiquitinated XBP1u was detectable in both ribosome and cytosol fractions in WT cells (Figure 3B). However, the depletion of LTN1 in LKO cells primarily reduced the ubiquitin signal in the ribosome fraction, supporting the notion that cytosolic ubiquitination of XBP1u is independent of LTN1 activity. Probing with a K48-linked ubiquitin chain-specific antibody revealed that nascent chains on ribosomes were modified with K48-linked ubiquitin chains by LTN1 (Figure S3A). Complementation experiment confirmed that the catalytic RING domain of LTN1 is essential for the ubiquitination of XBP1u nascent chains on stalled ribosomes (Figure S3B). In contrast to XBP1u, we observed only monoubiquitinated XBP1u-ΔHR2 in the cytosol fractions of both WT and LKO cells (Figure 3C). Since XBP1u-ΔHR2 contains the C-terminus AP, it demonstrated LTN1-dependent ubiquitination in the ribosome fractions. As anticipated, the stalling mutant XBP1u-AP* was exclusively ubiquitinated in the cytosol, and this ubiquitination was similar in both WT and LKO cells (Figure 3D).

**Figure 3.**
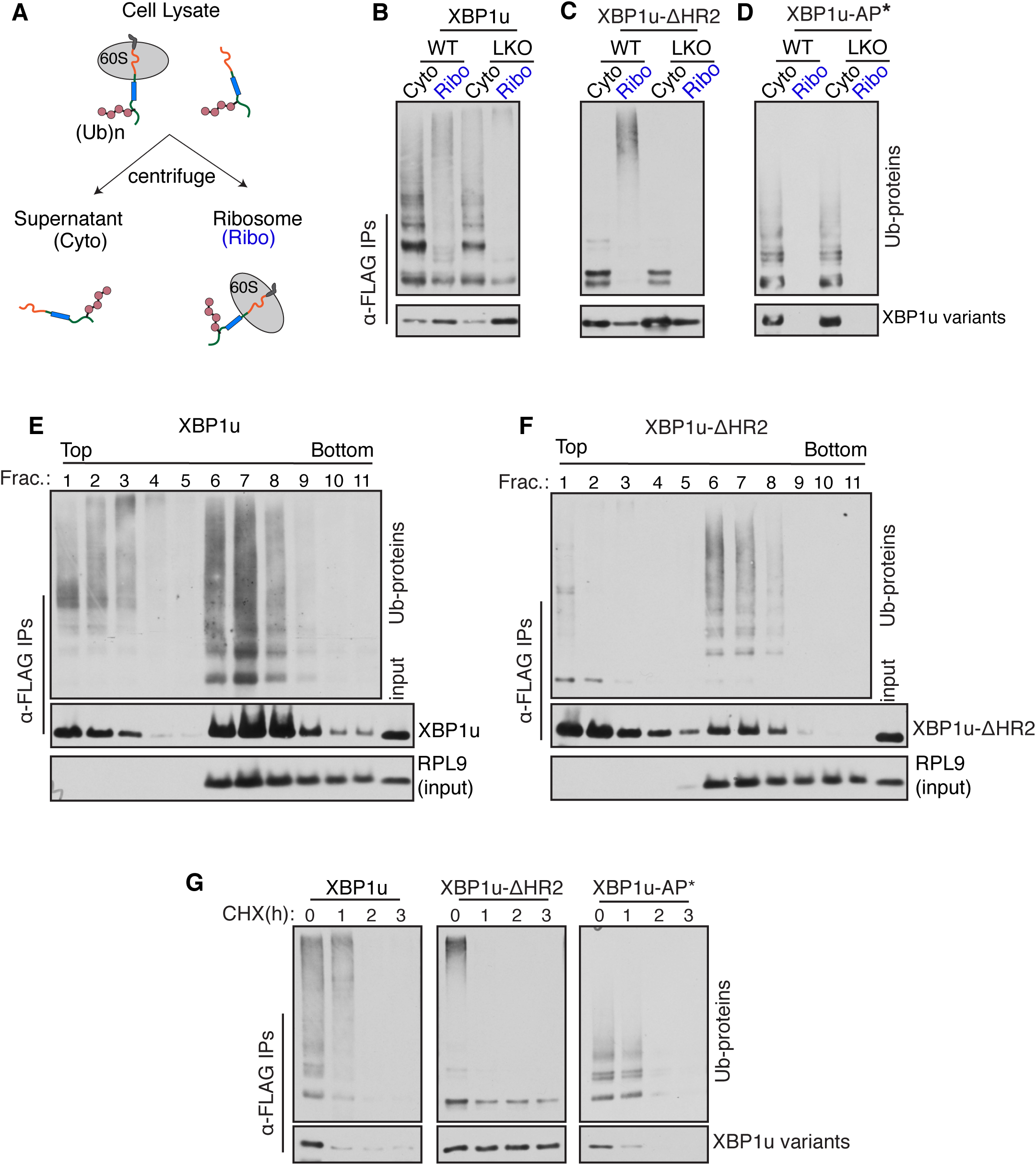
Nascent chains ubiquitinated by LTN1 are deubiquitinated in the cytosol. (A) Schematic depicting the separation of ribosome-associated nascent chains from free nascent chains in the cytosol through ultracentrifugation. (B, C, D) WT and LKO cells expressing the indicated XBP1u variant, along with HA-ubiquitin, were separated into cytosol and ribosome fractions. The resulting samples were immunoprecipitated using anti-FLAG beads and probed for Ub-proteins using an HA antibody. The presence of XBP1u variants was probed with an anti-FLAG antibody. (E, F) Cell lysates of XBP1u or XBP1uDHR2 were subjected to a 10-50% sucrose density gradient. Input fractions were directly probed for the ribosomal protein RPL9, while XBP1u and its mutant were immunoprecipitated using anti-FLAG beads and analyzed by immunoblotting. (G). Cells expressing the specified variant of XBP1u alongside HA-ubiquitin were treated with cycloheximide and collected at the indicated time points. The samples were subjected to anti-FLAG IP and were probed for Ub-proteins using an HA antibody. XBPu and its mutants were detected using anti-FLAG antibodies. See also **Figure S3.**

To further support the aforementioned observation that XBP1u undergoes two rounds of ubiquitination, we fractionated cell lysates on a sucrose density gradient. Immunoprecipitation for XBP1u was performed on all fractions, followed by probing for ubiquitinated proteins. Immunoblotting for ribosomal protein RPL9 indicated that the majority of ribosomes localized in fractions 6 to 8 (Figure 3E). XBP1u exhibited both AP- and HR2-dependent ubiquitination, as evidenced by the presence of ubiquitinated bands in both the peak ribosomal fractions (6-8) and the non-ribosomal fractions (1-3) (Figure 3E). In sharp contrast, XBP1u-ΔHR2 displayed AP-dependent ubiquitination in the ribosomal fractions but largely lacked HR2-dependent ubiquitin modifications in the non-ribosomal fractions (Figure 3F). AP-dependent ubiquitination for both XBP1u and XBP1u-ΔHR2 observed in the ribosomal fractions was diminished in LKO cells, further confirming that LTN1 is responsible for the ubiquitination of nascent chains on stalled ribosomes (Figure S3C and S3D). These results lead to the hypothesis that AP-dependent ubiquitin modifications are removed from nascent chains. To test this hypothesis, we monitored the status of ubiquitin modifications on XBP1u after inhibiting protein synthesis with cycloheximide (CHX). Both ubiquitin and total protein signals diminished during the CHX chase, indicating that ubiquitinated XBP1u is rapidly degraded (Figure 3G). In contrast, ubiquitin chains of ΔHR2, which occurred only on ribosomes (Figure 3F), were swiftly removed with little change in the ΔHR2 protein levels during a 3-hour chase (Figure 3G), indicating that AP-dependent ubiquitin chains undergo deubiquitination. The disappearance of ubiquitin signals for XBP1-AP*, along with a concomitant loss of total proteins, supports the notion that HR2-dependent ubiquitination is sufficient for proteasomal degradation (Figure 3G).

### Dissociation of nascent chains from stalled ribosomes results in deubiquitination in the cytosol

To elucidate the mechanism of XBP1u nascent chain deubiquitination, we reconstituted ubiquitination and deubiquitination reactions in vitro. Given that XBP1u AP-induced translational stalling is known to be transient in vitro^26^, we generated ribosome-associated nascent chains (RNCs) of XBP1u by translating transcripts lacking a stop codon in the rabbit reticulocyte lysate (RRL)-based translation system, including His-ubiquitin and radiolabeled 35S-methionine. The translationally stalled RNCs were ubiquitinated in the RRL, as evidenced by talon pull-down, which enriched for His-tagged ubiquitin-conjugated XBP1u or XBP1u-ΔHR2 (Figure 4A). Furthermore, the ubiquitination of RNCs was notably enhanced when incubated with cytosol prepared from WT HeLa cells. In contrast, incubation with cytosol derived from LKO cells did not significantly increase ubiquitination (Figure 4A). This result confirms that LTN1 E3 ligase mediates the ubiquitination of nascent chains on stalled ribosomes in vitro. Notably, the level of XBP1u-ΔHR2 ubiquitination was comparable to WT XBP1u, indicating that LTN1 mediates the ubiquitination of nascent chains on stalled ribosomes regardless of whether they contain exposed hydrophobicity (HR2). To determine if nascent chain dissociation from ribosomes alone is sufficient for deubiquitination, we treated RNCs with puromycin in the presence of cytosol. Puromycin mimics aminoacyl tRNA and releases nascent chains as peptidyl-puromycin derivatives^53^. Puromycin treatment of XBP1u RNCs resulted in the loss of polyubiquitinated species and the detection of lower ubiquitinated species (Figure 4B). These bands did not appear when an E1 inhibitor (MLN4924)^54^, which inhibits re-ubiquitination, was included during the puromycin treatment. This suggests that they represent re-ubiquitination of XBP1u in the cytosol rather than leftover ubiquitin modifications from deubiquitination. In contrast, the ubiquitin chains of XBP1u-ΔHR2 were nearly entirely lost upon puromycin treatment, with little re-ubiquitination occurring (Figure 4B). Fractionation by sucrose cushion centrifugation after puromycin treatment further revealed that re-ubiquitinated XBP1u and deubiquitinated XBP1u-ΔHR2 were localized to the supernatant cytosol fractions (Figure 4C).

**Figure 4.**
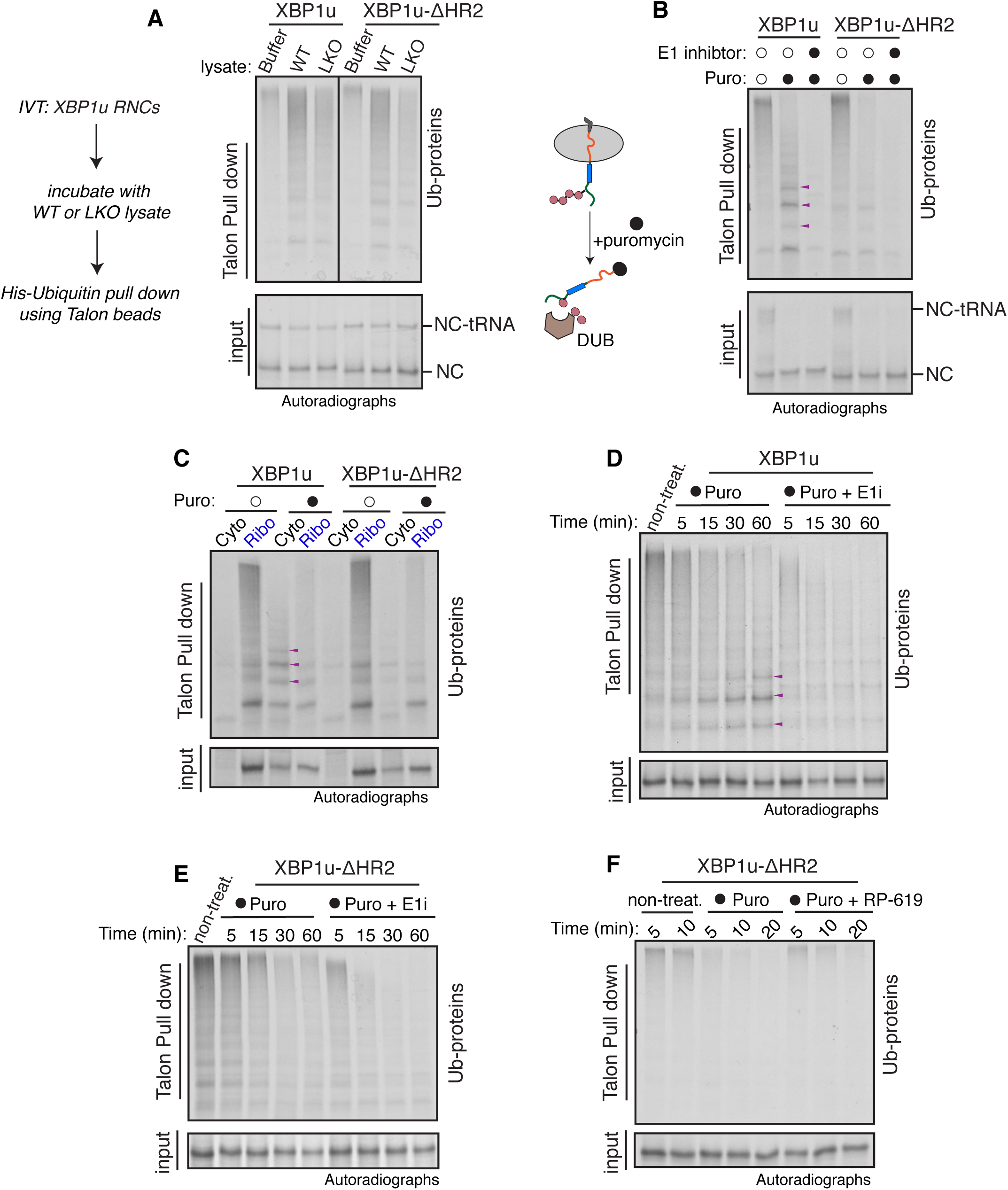
Nascent chain is deubiquitinated after its dissociation from the ribosome. (A) The indicated XBP1u transcripts lacking a termination codon were translated in rabbit reticulocyte lysate (RRL) supplemented with 35S-methionine and His-ubiquitin. The reactions containing ribosome-associated nascent chains (RNCs) were incubated with a buffer or the cytosol from WT HeLa or LKO cells. The samples were then denatured and either directly analyzed as input or subjected to Talon pull-down to capture His-ubiquitin-conjugated radiolabeled XBP1u. The resulting samples were visualized by autoradiography. Notably, tRNA-attached nascent chains (NC) were preserved during SDS/PAGE electrophoresis. (B) Ubiquitinated RNCs of XBP1u and its mutant were prepared as in (A) and subsequently treated with puromycin for 45 minutes. An E1 inhibitor (MLN4924) was included along with puromycin. The samples were then denatured and analyzed as in (A). Arrowhead indicates the re-ubiquitinated bands. (C) The reactions were performed as described in (B), with subsequent analysis conducted after separating cytosol from ribosomes through ultracentrifugation. The Arrowhead indicates the re-ubiquitinated bands. (D, E) Ubiquitinated RNCs were prepared and analyzed as described in (B), with samples collected at various time points during the incubation for subsequent analysis. Arrowhead in (D) indicates the re-ubiquitinated bands. (F) Ubiquitinated RNCs of XBP1uDHR2 were prepared and analyzed as in (B), with the addition of a generic DUB inhibitor PR-619 in one set of reactions during puromycin treatment. See also **Figure S4.**

Next, we monitored the dynamics of deubiquitination and re-ubiquitination by treating RNCs with puromycin in the presence of cytosol and sampled at various time points to assess ubiquitin-conjugated proteins. Figure 4D shows that polyubiquitinated species mostly disappeared within 15 minutes of nascent chain release by puromycin. Remarkably, newly ubiquitinated XBP1u emerged after 30 minutes, indicating that re-ubiquitination occurs later. The re-ubiquitination of XBP1u could be entirely inhibited when the E1 inhibitor was included along with puromycin, providing support for the involvement of a cytosolic E3 ligase(s) in mediating the re-ubiquitination of XBP1u (Figure 4D). In contrast to XBP1u, ubiquitin chains on XBP1u-ΔHR2 were largely removed within 30 minutes with no new ubiquitin modifications occurring, and little change was observed when including the E1 inhibitor (Figure 4E). This result implies that the exposed hydrophobicity of HR2 in XBP1u is strictly required for re-ubiquitination in the cytosol.

Cytosolic deubiquitinases (DUBs) were responsible for removing ubiquitin chains from nascent chains, as evidenced by the reduced deubiquitination of nascent chains when the cytosol was treated with the generic DUB inhibitor PR-619^55^ (Figure 4F). These findings suggest that the dissociation of nascent chains from the ribosome-LTN1 E3 ligase complex is sufficient for their deubiquitination by cytosolic DUBs. Additionally, our results propose a paradigm whereby substrates undergo continuous modifications by their respective E3 ligases to effectively compete against cytosolic DUBs. To explore this, we generated polyubiquitinated XBP1u or XBP1u-ΔHR2 RNCs and incubated them with untreated or E1 inhibitor-treated cytosol. Strikingly, ubiquitin chains on nascent chains were mostly removed in the presence of E1 inhibitor-treated cytosol, even though they remained attached to stalled ribosomes (Figure S4A). This observation is not exclusive to LTN1-mediated ubiquitination occurring on stalled ribosomes, as the stalling mutant XBP1u-AP*, ubiquitinated by the cytosolic quality control, also underwent deubiquitination when incubated with the E1 inhibitor (Figure S4B). Collectively, these data suggest a model where DUBs constantly remove ubiquitin chains from nascent chains, providing an opportunity for protein maturation. However, if a deubiquitinated protein consistently exposes a hydrophobic region, as observed with XBP1u, it is then recognized by the cytosolic quality control for re-ubiquitination and degradation.

### Protein folding status dictates stability versus re-ubiquitination of deubiquitinated nascent proteins

To further support our model, which suggests that cytosolic quality control, rather than ubiquitination occurring on ribosomes, governs the degradation of nascent chains, we aimed to identify the quality control machinery responsible for recognizing the exposed hydrophobicity (HR2) in XBP1u in the cytosol. Given the established role of the Bag6 complex in recognizing long hydrophobic sequences in proteins and mediating ubiquitination in the cytosol^24,56,57^, we performed a co-immunoprecipitation to probe the interaction between XBP1u and Bag6. Bag6 interreacted with XBP1u and XBP1u-AP*, both of which contain HR2, and weakly interacted with XBP1u-ΔHR2, as expected (Figure 5A). Notably, XBP1u interaction with Bag6 was further intensified when proteasomes were inhibited with MG132, indicating that Bag6-associated XBP1u is routed for proteasomal degradation. Indeed, siRNA-mediated depletion of Bag6 led to the accumulation of XBP1u in the cytosol fraction compared to cells treated with control siRNA (Figure 5B). We observed that Bag6 depletion led to partial inhibition of cell growth, consequently resulting in reduced expression of XBP1u in these cells compared to those treated with control siRNA. Furthermore, effective XBP1u re-ubiquitination in the cytosol depends on its binding with Bag6, as its ubiquitination was significantly reduced in the cytosol fraction of Bag6-depleted cells compared to cells treated with control siRNA (Figure 5C). To investigate whether the Bag6-associated E3 ligase, RNF126^58^, is involved in XBP1u degradation, we depleted RNF126 using siRNA and monitored XBP1u ubiquitination and degradation. Indeed, both ubiquitination and degradation of XBP1u were dependent on the activity of RNF126 E3 ligase, as evidenced by reduced ubiquitination and impaired degradation in RNF126-depleted cells compared to control siRNA treated cells (Figure 5D, 5E, and Figure S5A).

**Figure 5.**
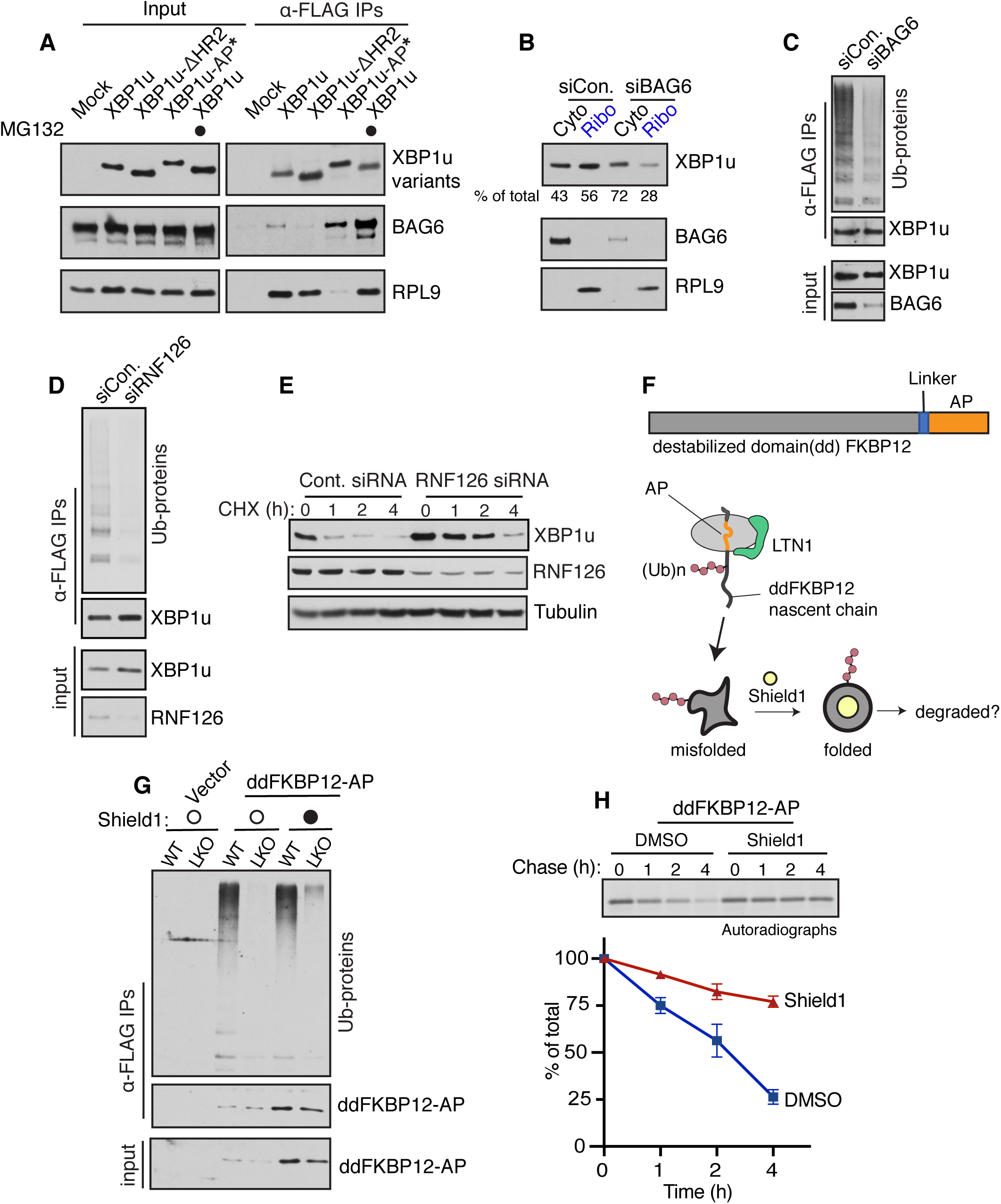
Exposed hydrophobicity determines re-ubiquitination and degradation of deubiquitinated nascent proteins. (A) Cells expressing the indicated HA-tagged constructs were subjected to anti-HA magnetic beads and analyzed by immunoblotting for the indicated antigens. (B) Cells treated with either control siRNA or Bag6 siRNA were transfected with FLAG-XBP1u and HA-ubiquitin, followed by fractionation into cytosolic and ribosomal fractions for subsequent analysis by immunoblotting. (C) The cytosol fraction from (B) was immunoprecipitated with anti-FLAG beads and analyzed by immunoblotting. (D) FLAG-XBP1u was transfected with HA-ubiquitin into cells treated with either control siRNA or RNF126 siRNA. Cell lysates were immunoprecipitated with anti-FLAG beads and analyzed by immunoblotting. (E) CHX chase analysis was performed in cells treated with indicated siRNA and expressing XBP1u, followed by immunoblotting for the indicated antigens. (F) (Top) Model construct illustrating the destabilized domain (dd) comprising FKBP12 fused with the arrest peptide (AP) of XBP1u. (Bottom) Schematic representation of LTN1-mediated ubiquitination of ddFKBP12-AP nascent chain. Shield1 small molecule is known to stabilize the folding of ddFKBP12. (G) WT or LKO cells expressing dd FKBP12-AP alongside HA-ubiquitin were untreated or treated with Shield1, immunoprecipitated, and analyzed by immunoblotting. (H) Pulse-chase experiments were conducted using cells expressing ddFKBP12-AP, treated either with or without Shield1. See also **Figure S5.**

Our data so far suggest that exposed hydrophobicity, often exhibited by misfolded proteins, is pivotal in driving re-ubiquitination of deubiquitinated nascent proteins in the cytosol, ultimately leading to proteasomal degradation. To scrutinize this model directly, we capitalized a misfolded form of FKBP12 containing an engineered destabilized domain (dd). This misfolded protein is constitutively degraded in cells, but the addition of Shield-1, which binds to the ddFKBP12 and stabilizes the protein fold, rescues them from degradation^59^. We hypothesized that if protein misfolding in the cytosol is the primary factor influencing stalled nascent chain degradation, misfolded ddFKBP12 should be degraded but stabilized in the presence of the fold stabilizer Shield1, even when ubiquitinated on stalled ribosomes by LTN1 (Figure 5F). To test this, the C-terminus of ddFKBP12 was appended with the AP sequence from XBP1u. ddFKBP-AP underwent ubiquitination in an LTN1-dependent manner, both in the presence and absence of the folding stabilizer Shield1 (Figure 5G). As expected, ddFKBP-AP was subject to degradation in the absence of Shield1 due to its misfolding (Figure 5H and Figure S5B). Surprisingly, ddFKBP-AP exhibited significant stabilization in the presence of the fold stabilizer Shield1 despite being ubiquitinated by LTN1 on stalled ribosomes. We obtained similar results when ddFKBP was fused with a stronger AP sequence, which is known to induce stronger translational pausing^26^ (Figure S5B and Figure S5C). These results provide further support for the model that the folding status determines whether a deubiquitinated nascent protein becomes stable or undergoes degradation in the cytosol, rather than the LTN1-mediated ubiquitination of nascent chains occurring on ribosomes.

### LTN1 E3 ligase activity is critical for eliciting proper UPR

Given that LTN1-mediated ubiquitination is found to be dispensable for proteasomal degradation, we asked what is the purpose of LTN1-mediated ubiquitination. We hypothesized that the activity of the LTN1 E3 ligase is crucial for releasing nascent chains from stalled ribosomes, thus allowing them to either mature or undergo degradation based on their folding status in the cytosol. To explore this hypothesis, we initially examined the localization of endogenous XBP1u in LKO cells. Fractionation experiments revealed a predominant accumulation of endogenous XBP1u with ribosomes in LKO cells compared to WT cells, supporting LTN1-dependent release of XBP1u from stalled ribosomes (Figure 6A). The release of XBP1u from stalled ribosomes is dependent on the ligase activity of LTN1 because the complementation of the catalytic RING domain mutant (ΔRING) of LTN1 did not prevent XBP1u accumulation in the ribosome fraction of LKO cells (Figure 6B). By contrast, complementation with WT LTN1 restored the release of XBP1u from ribosomes (Figure 6B). We further investigated the LTN1-dependent release of XBP1u from stalled ribosomes by conducting localization studies in 293T cells. Immunofluorescence staining demonstrated that XBP1u primarily exhibits an ER-like pattern in both WT and LKO cells (Figure 6C). While only a small fraction of XBP1u was localized to the nucleus in non-treated cells, a brief inhibition of proteasomes with MG132 resulted in a significant accumulation of XBP1u in the nucleus of WT cells due to its nuclear localization signal^51,60^. In contrast, the localization pattern of XBP1u did not noticeably change upon MG132 treatment of LKO cells, as it remained confined to the ER (Figure 6C and 6D), supporting the LTN1-mediates the release of XBP1u nascent chains from the ER membrane. XBP1u was associated with the ER membrane as RNCs since it could be readily released by treating cells with puromycin prior to immunostaining (Figure S6). Furthermore, neutral page (NUPAGE)-based immunoblotting, which preserves the ester bonds between tRNAs and nascent chains during SDS-PAGE electrophoresis, revealed a notable accumulation of tRNA attached XBP1u nascent chains in LKO cells relative to WT cells. As anticipated, XBP1u-AP* showed a loss of tRNA-linked XBP1u, while the arrest enhancing mutant S255A^26^ showed increased levels of tRNA-linked XBP1u in WT cells, with further accumulation in LKO cells (Figure 6E).

**Figure 6.**
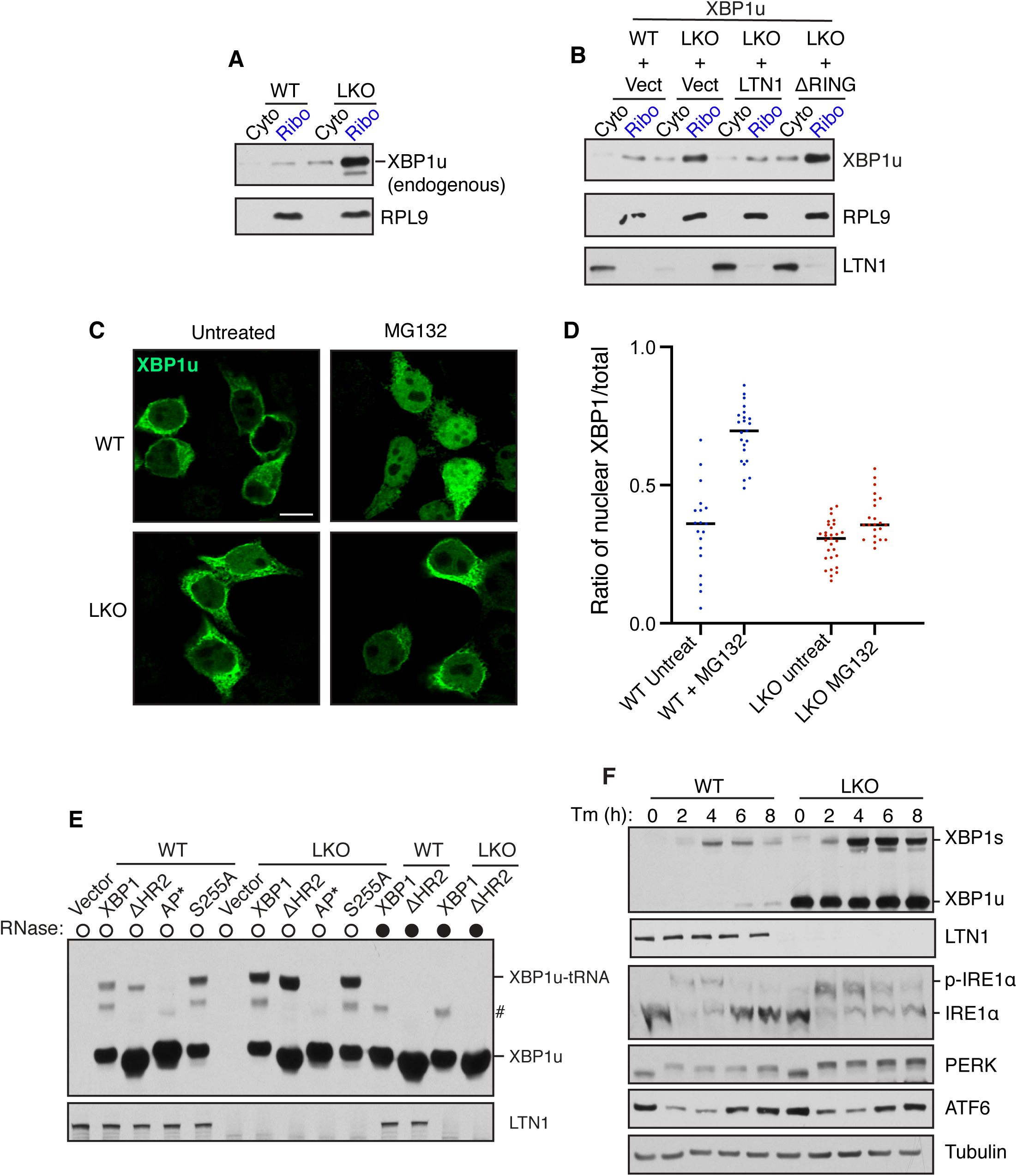
The unfolded protein response (UPR) is dysregulated in LTN1-depleted cells. (A) Immunoblotting analysis of endogenous XBP1u in supernatants and ribosome fractions from WT or LKO cells. (B) Cells expressing indicated plasmids were fractionated into supernatants and ribosomes and analyzed by immunoblotting for the indicated antigens. (C) WT or LKO cells expressing XBP1u were either untreated or treated with 20μM MG132 for 30min. Subsequently, cells were processed using an immunostaining procedure to label XBP1u (green) with rat anti-FLAG. Scale bar is 10 μm. (D) Quantification of the ratio of nuclear-localized XBP1u to the total in individual cells from panel (C). n> 19 from two independent experiments. (E) Cell lysates expressing the indicated variants of XBP1u were separated under neutral pH conditions (NuPAGE) to preserve peptidyl-tRNA ester bonds. Note that tRNA-linked XBP1u was sensitive to RNase treatment. # denotes an unknown band. (E) WT or LKO cells were treated with tunicamycin (Tm: 5μg/ml) and analyzed for immunoblotting for the indicated antigen. Note that the loss of signal represents the activation of ATF6 during ER stress. See also **Figure S6.**

Given the profound accumulation of XBP1u RNCs in the ER membrane of LKO cells, we wondered whether this would impact IRE1α-mediated XBP1u splicing during ER stress. Remarkably, the production of both spliced and unspliced forms of XBP1 was significantly increased in LKO cells treated with the ER stress inducer tunicamycin compared to WT cells (Figure 6F). Additionally, IRE1α exhibited hyperactivation and displayed a delay in inactivation during prolonged ER stress, as evidenced by the phosphorylation status of IRE1α probed by Phos-tag based immunoblotting (Figure 6F). Notably, the depletion of LTN1 selectively dysregulated the IRE1α branch of the UPR since the activation and inactivation of the ATF6α and PERK branches behaved similarly in both WT and LKO cells. This is consistent with our earlier findings that only the IRE1α branch is associated with the Sec61 translocon complex^50,61^, where XBP1u RNCs are recruited. Collectively, our data suggest that the central role of LTN1-mediated ubiquitination is to facilitate the release of nascent chains from stalled ribosomes, thereby contributing to maintaining protein homeostasis by recycling ribosomes.

## DISCUSSION

About 5% to 15% of nascent proteins undergo ubiquitination while on ribosomes^4,5^, but the mechanisms and features in these proteins that mediate proteasomal degradation have remained unclear. In this study, we systematically unraveled the degradation mechanisms of nascent chains ubiquitinated on ribosomes using a physiologically crucial protein, XBP1u, which induces ribosomal stalling through its C-terminal arrest peptide (AP) during translation elongation^26^. Our investigations revealed robust ubiquitination of the XBP1u nascent chain on stalled ribosomes by LTN1 E3 ligase. Strikingly, despite possessing K48-linked ubiquitin chains, which are typically associated with proteasomal degradation^62^, ubiquitin chains on nascent chains are promptly removed by cytosolic deubiquitinases upon their release from stalled ribosomes. Our data from in vitro experiments suggest that once the ubiquitinated nascent chain is liberated from the LTN1 E3 ligase-ribosome complex, it can no longer receive new ubiquitin modifications to effectively compete against cytosolic deubiquitinases, resulting in complete deubiquitination. By studying XBP1u mutants, we further found that deubiquitinated nascent proteins can attain stability (Figure 7). However, if they persistently expose a hydrophobic sequence, they become targets for recognition by the cytosolic quality control machinery. This leads to re-ubiquitination and subsequent proteasomal degradation.

**Figure 7:**
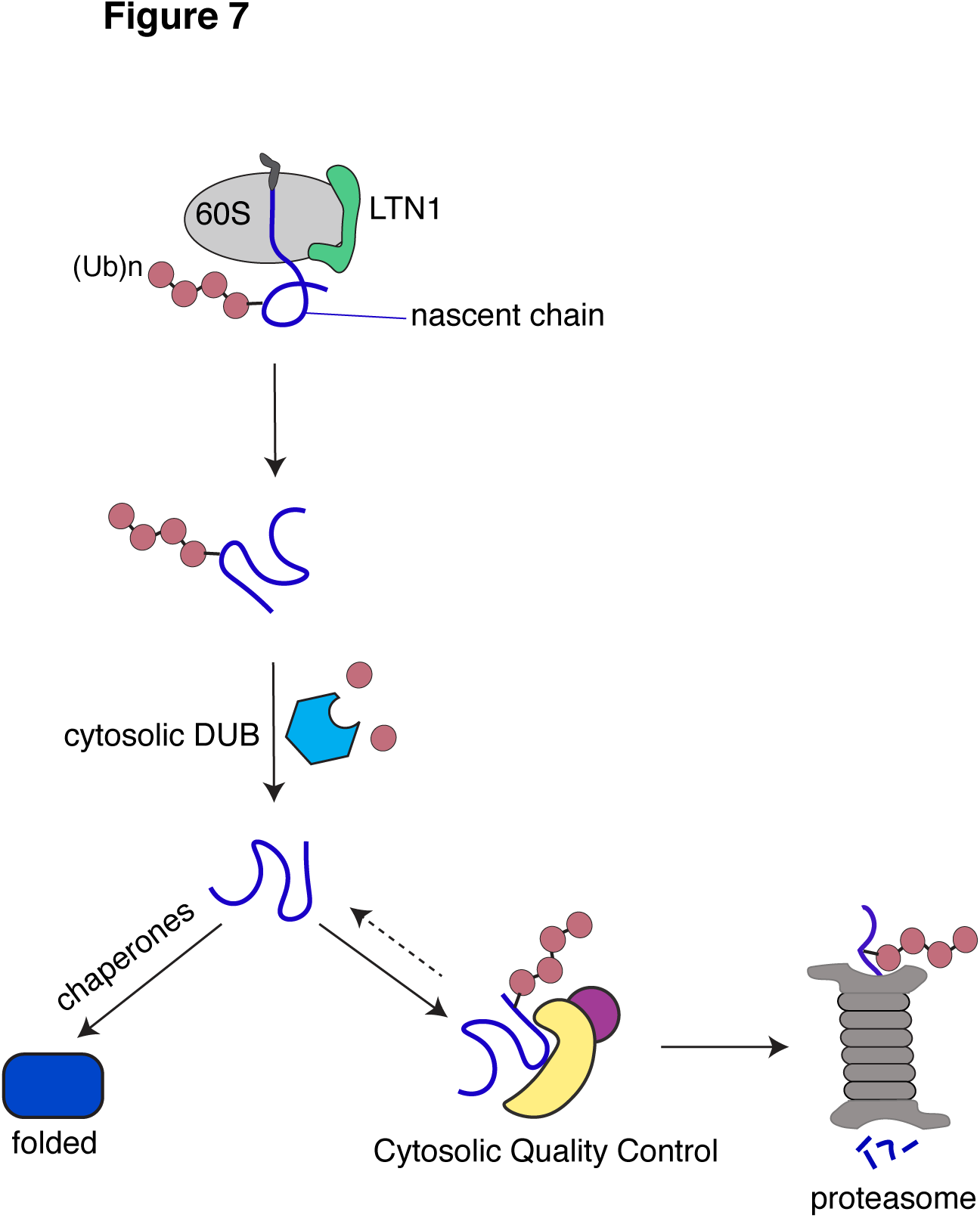
A model for triaging ubiquitinated nascent proteins between maturation and degradation. Upon release from stalled ribosomes, nascent proteins polyubiquitinated by the LTN1 E3 ligase are promptly deubiquitinated in the cytosol. This decoupling from immediate proteasomal degradation provides a secondary chance for some nascent proteins to achieve proper folding and maturation. However, if a nascent protein fails to attain stability and exposes a degron, it undergoes re-ubiquitination through cytosolic quality control mechanisms, leading to proteasomal degradation. It is conceivable that the nascent protein may re-enter a similar cycle, but ultimately, it would face degradation by the proteasome if it remains exposed to a degron for an extended period.

The data presented here highlight that the nascent chain’s biophysical properties, particularly its folding status as supported by the results using a conditionally folded FKBP12 mutant (Figure 5G, 5H), play a crucial role in determining whether it engages with the cytosolic quality control for degradation or achieves stability. This is in contrast to the initial ubiquitination event occurring on ribosomes, which does not seem to be the primary determinant of the nascent chain’s fate. The mechanism revealed in our studies acts as a safeguard, potentially preventing the indiscriminate degradation of numerous proteins known to induce ribosomal stalling through their C-terminal sequences^20,21^. This enables these proteins to potentially fold into functional forms in the cytosol, even if they were initially ubiquitinated on stalled ribosomes.

Our findings may also shed light on why cells have evolved an alternative but conserved mechanism to add C-terminal Ala tails in mammals^63^ (or C-terminal Ala and Thr tracts known as CAT tails in yeast,)^36,64^ in an mRNA-independent but NEMF-assisted manner. This C-terminal tail ensures the recruitment of the cytosolic quality control for ubiquitination and subsequent degradation of stalled nascent proteins^63,65^. It appears unlikely that XBP1u undergoes NEMF-mediated mRNA-independent C-terminal alanine tailing. This is because the translational elongation facilitated by NEMF may not be able to advance the XBP1u AP, which interacts with components of the ribosome exit tunnel^49^. It is tempting to speculate that commonly used C-terminal poly lysines containing RQC substrates may function as a degron recognized by a cytosolic quality E3 ligase for a second round of ubiquitination and subsequent proteasomal degradation. This would explain the discrepancy between our results, which show complete stabilization of ddFKBP12-AP in the presence of the fold stabilizer Shield 1, and earlier reports demonstrating the degradation of ddFKBP12 fused with a poly(A) tail (encoding poly lysines)^43^. By contrast, the AP of XBP1u does not have any obvious degron motif for recognition by cytosolic quality control.

While our results demonstrate that LTN1-mediated ubiquitination of nascent chains is not essential for directing them toward proteasomal degradation, the activity of the LTN1 E3 ligase is required for the efficient release of XBP1u nascent chains from stalled ribosomes (Figure 6). Failure to release XBP1u RNCs from the ER membrane in LKO cells leads to heightened activation of IRE1α and elevated levels of spliced XBP1 factor during ER stress. Further studies are warranted to precisely elucidate how LTN1-mediated ubiquitination facilitates the release of nascent chains from stalled ribosomes bound to the Sec61 translocon. Additionally, it will be important to understand the coordination between LTN1-mediated release of XBP1u and IRE1α-mediated splicing of XBP1u during ER stress.

One notable observation from biochemical reconstitution experiments is the complete deubiquitination of XBP1u following a brief treatment with an E1 inhibitor (Figure S4). This suggests that substrates must undergo continuous ubiquitination to effectively compete with cytosolic DUBs, which is consistent with the earlier observation that cytosolic DUBs can efficiently trim ubiquitin chains from the substrate that interacts weakly with an E3 ligase^66^. Our results also imply that a substrate should consistently present a degron, allowing it to engage with an E3 ligase complex for efficient ubiquitination and subsequent degradation by proteasomes.

The presented data suggest that the re-ubiquitination of stalled nascent chains in the cytosol is an essential step for their subsequent proteasomal degradation (Figure 7). Yet, it remains a critical area of investigation whether these nascent chains, ubiquitinated for the second time in the cytosol, are directly shuttled for proteasomal degradation. Alternatively, it is plausible that these proteins undergo cycles of deubiquitination and re-ubiquitination before ultimately reaching the degradation stage. This mechanism is similar to how N-glycans play a crucial role in overseeing the folding and quality control of nascent glycoproteins^67^. This cycle of ubiquitination and deubiquitination may explain why proteasomal degradation of a substrate is very slow and often takes hours in cells, which is in contrast to the relatively rapid ubiquitination and proteasomal degradation that can occur within minutes for proteins of average size^68,69^.

In summary, our studies suggest that cells have uncoupled ubiquitination from proteasomal degradation to provide chances for the maturation of folding-competent proteins while ensuring the degradation of terminally misfolded proteins that consistently expose a degron. However, this mechanism poses a risk of accumulating aberrant nascent proteins that cannot display degron signatures in the cytosol. For instance, deubiquitination of disease-causing proteins like huntingtin (mHTT), which is known to induce ribosome stalling during translation^70,71^, could delay proteasomal degradation and promote aggregation of mHTT in the cytosol. Moreover, during proteotoxic stress conditions such as aging, re-ubiquitination of stalled aberrant nascent proteins may be inefficient in the cytosol, potentially leading to the accumulation of toxic proteins in cells. Hence, targeting cytosolic DUBs responsible for deubiquitinating nascent chains could hold therapeutic potential for promoting the degradation of mHTT and other stalled aberrant nascent proteins, which are particularly exacerbated during aging^23^.

### Limitations of the study

While we have shown the recruitment of NEMF and LTN1 to XBP1u nascent chains on stalled ribosomes, our studies do not definitively establish whether LTN1 binds to 60S subunits and mediates the ubiquitination of XBP1u nascent chains attached to them. Although our investigation into the endogenous staller XBP1u rigorously demonstrates that LTN1-mediated ubiquitination occurring on ribosomes is insufficient to mediate proteasomal degradation, further work is needed to identify additional endogenous substrates ubiquitinated on ribosomes and to assess if they follow similar regulatory principles. Both our in vitro and in vivo experiments suggest that nascent chains ubiquitinated on ribosomes are deubiquitinated as soon as they dissociate from ribosomes. However, future studies are warranted to identify specific DUBs involved in deubiquitinating nascent chains in the cytosol.

## ACKNOWLEDGEMENTS

We want to thank the former and present members of the Mariappan lab for valuable discussions. We thank Jean Kanyo and TuKiet Lam from the Yale Keck Proteomic facility for assisting with MS data. We are grateful to Susan Shao for providing the LTN1 and LTN1 (ΔRING) plasmids and to Zairong Zhang for the ddFKBP12 plasmid. This work was supported by NIH grant R01GM117386 (M.M) and a pilot award from the Yale Diabetes Research Center (P30 DK045735).

## AUTHOR CONTRIBUTIONS

XL designed and conducted all experiments. MM conceived and oversaw the project. MM drafted the manuscript with contributions from XL.

## STAR*METHODS

### RESOURCE AVAILABILITY

#### Lead Contact

Further information and requests for resources and reagents should be directed to and will be fulfilled by the Lead Contact, Malaiyalam Mariappan.

#### Materials Availability

All reagents generated in this study are available from the corresponding author with a completed Materials Transfer Agreement.

#### Data and Code Availability

Original data will be made available upon request to the lead contact.

### EXPERIMENTAL MODEL AND SUBJECT DETAILS

HEK 293T, HeLa, and derivative cells were cultured in Dulbecco’s Modified Eagle’s Medium (DMEM) and 10% fetal bovine serum (FBS) and 100 U/mL penicillin and 100 µg/mL streptomycin at 5% CO2. LTN1 knockout (LKO) cells created by CRISPR/Cas9 were previously described in method details. Knock-out and stable cell lines were routinely verified by immunoblotting.

### METHOD DETAILS

#### DNA constructs

The mammalian cell expression vector pcDNA5/FRT/TO (Invitrogen, Carlsbad, CA) containing FLAG-tagged XBP1u, XBP1u-ΔHR2, XBP1u-AP* (L246A and W256A), or XBP1u (S255A) was previously detailed^50^. XBP1u-4L was created by introducing Q197L, Q198L, S200L, and S203L mutations using a site-directed mutagenesis protocol. Mammalian expression constructs of FLAG-LTN1 and FLAG-LTN1ΔRING were kindly provided by Dr. Susan Shao (Harvard Medical School). FLAG-XBP1u-step was constructed by replacing the HR2 with a strep-tag sequence (SAWSHPQFERGGSWSHPQFERAS). XBP1u arrest peptide (AP) sequence of 26 amino acids was fused to the C-terminus of the destabilizing mutant of FKBP12 L106P (kindly provided by Dr. Zairong Zhang, University of Chinese Academy of Sciences) using a standard cloning procedure. All PCR reactions were performed with Phusion high-fidelity DNA polymerase (New England Biolabs, Ipswich, MA), except for site-directed mutagenesis, which used Pfu-Ultra polymerase (Agilent Technologies, Santa Clara, CA). 3% DMSO was included in all PCR reactions to enhance amplification. The coding regions of all constructs were sequenced to prevent any sequence errors.

#### Cell culture and generation of CRISPR/Cas9-mediated knockout cell lines

HEK293T and HeLa cells were cultured in high-glucose DMEM (Corning) supplemented with 10% FBS (Gibco), 100 U/ml penicillin, and 100 mg/ml streptomycin (Gibco) at 37°C with 5% CO2. Regular mycoplasma testing was performed using the MycoStrip kit from Invivogen. LTN1 siRNA (CAAACUUCCUGCAAGAUUA), RNF126 siRNA (CGGAUUAUAUCUGUCCAAG), Bag6 siRNA (GGACAAACCUGGAAUUUCU) were synthesized from Sigma-Aldrich. Knockdown was performed using Lipofectamine RNAiMAX (Invitrogen) according to the manufacture protocol. Knockout cell lines were generated using the CRISPR/Cas9 system as previously described^72,73^. Cells were transfected with pSpCas9(BB)-2A-Puro (Addgene; #62988) and gRNA expression plasmid targeting LTN1. Two targeting sequences were used: 5′-AATGCCGAACGACAGTGAA-3′ and 5′-CGTGTACTTGGTAATACGT-3′. gRNAs were designed using the web tool CHOPCHOP (version 3)^74^. Transfected cells were grown for 24 hours and then treated with 4µg/ml puromycin for 3 to 4 days until all control cells had died. Most experiments were conducted using mixed knockout clones of LTN1, as these exhibited over 90% depletion of LTN1, as confirmed by immunoblotting. In some cases, single-cell clones were isolated by plating at a density of 0.5 cells/well in 96-well plates. All knockouts were confirmed by immunoblotting.

#### Affinity purification of XBP1u and mass spectrometry

To identify interacting proteins of XBP1u through mass spectrometry, 293T cells were seeded into five 150 mm dishes at a density of 5 x 10^6^ cells per dish. The following day, cells were transfected with either 10 μg of control or Flag-XBP1u plasmid using 40 μl of PEI reagent. After 36 hours of transfection, cells were harvested in cold PBS and centrifuged at 5,000 g for 5 minutes. Cell pellets were then lysed in 5 ml of ice-cold lysis buffer (50 mM Hepes, 125 mM KCL, 10 mM MgCl2, 0.1% Triton, 1 mM DTT, 20 U/ml RNase inhibitor, 20 μM MG132, 2 mM NEM, protease inhibitor). The clear supernatant of the cell lysate was obtained after two rounds of centrifugation (700 g for 10 minutes, followed by 20,000 g for 15 minutes in a cold room), and mixed with 100 μl of compact FLAG-M2 beads (previously rinsed with lysis buffer) for 1.5 hours. The beads were subsequently extensively washed with lysis buffer.

For protein elution, 100 μl of FLAG peptide elution buffer (0.2 mg/ml of 3xFLAG peptide (Sigma), 50 mM Hepes, 125 mM KCl, 10 mM MgCl2, 1 mM DTT, 0.1% Triton) was added to the beads. After incubating at room temperature for 20 minutes, the mixture was centrifuged at 5,000 rpm for 3 minutes to collect the eluted protein supernatant. The elution step was repeated twice, pooling the eluted proteins. The beads were then resuspended with 500 μl of elution buffer without FLAG peptide and centrifuged to collect the supernatant containing the eluted proteins. The eluate was passed through a 1 ml Bio-rad column to remove residual beads. The eluted materials were TCA precipitated and subjected to mass spectrometry analysis at the Yale Keck Proteomic facility. Statistical analysis of the results utilized the total spectrum count of each protein from the control or FLAG-XBP1u immunoprecipitation (IP). A scatter plot was generated, plotting the log2 of the total spectrum count for each protein.

#### Metabolic labeling and immunoprecipitation

HEK293T cells (0.35 × 10^6^/well) were seeded onto polylysine-coated plates (0.15 mg/ml) in 12-well format and transiently transfected with 0.8 µg of the specified plasmids using 2 µl of Lipofectamine 2000 (Invitrogen). For metabolic labeling, cells were washed once with 1× PBS and then incubated in cysteine- and methionine-free media supplemented with 5% dialyzed FBS for 45 minutes. Subsequently, cells were labeled with 80 µCi/ml of Express ^35^S Protein Labeling Mix (PerkinElmer; #NEG072014MC) for 30 minutes. The cells were then rinsed with 1× PBS and chased with complete DMEM medium containing 2 mM methionine and 2 mM cysteine. The labeled cells were directly lysed in 0.5 ml RIPA buffer and centrifuged for 15 minutes at 20,000xg. The resulting supernatant was subjected to immunoprecipitation using rat anti-FLAG beads (BioLegend). The immunoprecipitants were subsequently analyzed by SDS-PAGE followed by autoradiography

#### Cycloheximide chase

HEK293T (0.125 × 10^6^/well) were plated on a polylysine pre-coated 24-well–plate and transiently transfected with 0.4 μg of the indicated construct as described in the figure legends. After 24h of transfection, cells were treated with cycloheximide (100 μg/ml). The treated cells were directly harvested with 150 μl of 2× SDS sample buffer at the indicated time points. In some cases, cells were treated with both cycloheximide and MG132 (20 μM). Samples were then analyzed by immunoblotting with the indicated antibodies as described in the figure legends.

#### Immunoprecipitation of ubiquitinated proteins from cells

To examine total ubiquitination of XBP1u and its mutants, HEK293 cells (0.35 × 10^6^/well) were seeded on polylysine-coated six-well plates and transiently transfected with 0.6 µg of XBP1u and 0.4 µg of HA-Ubiquitin using 2 µl of Lipofectamine-2000. After 24 hours of transfection, cells were washed once with 1xPBS and directly harvested using 120 µl of Tris/SDS buffer (1% SDS/100mM Tris-HCl pH 8.0). The samples were boiled for 10 minutes and a 15 µl aliquot was directly analyzed by immunoblotting as input. The remaining samples were diluted with 1 ml of Triton buffer (50mM Tris-HCl pH 8.0, 150mM NaCl, 1% Triton X-100) and immunoprecipitated using anti-FLAG beads, followed by immunoblotting analysis.

#### Sucrose density gradient analysis

For the sucrose gradient analysis presented in Figure 1C, HEK293T cells (1.5 × 10^6^/well) were seeded onto polylysine-coated 60 cm dishes and transiently transfected with 4µg of the specified plasmids using 16 µl of PEI (1mg/ml). After 24 hours of transfection, cells were harvested in 1x PBS. The cell pellet was resuspended in 250 µl of lysis buffer (50mM Hepes, pH 7.4, 100mM KAc, 5mM MgCl2, 1% Triton X-100, 2.5u/ml of RNasin (Promega), and EDTA-free protease inhibitor (PI) from Roche) and incubated for 15 minutes on ice. Cell lysates were centrifuged at 20,000xg for 5 minutes. The resulting supernatant (200 µl) was carefully applied onto a 10% to 30% sucrose gradient and centrifuged for 80 minutes at 215,000xg using a TLS-55 rotor at 4°C with the slowest acceleration and deceleration settings. After the centrifugation, 12 x 165 µl fractions were collected and subsequently analyzed by immunoblotting.

For the sucrose gradient data presented in Figure 3E, 3F, S3B, and S3C, HEK293T cells (4 × 10^6^/well) were seeded onto polylysine-coated 10 cm dishes and transiently transfected with 8µg of FLAG-XBP1u and 4µg of HA-Ubiquitin using 40 µl of PEI (1mg/ml). After 24 hours of transfection, cells were harvested in ice-cold 1x PBS. The cell pellet was resuspended in 300 µl of ribosome buffer (50mM Hepes, pH 7.4, 500mM KCl, 10mM MgCl2, 0.5% Triton X-100, 2.5u/ml of RNasin (Promega), and 1x PI) and incubated for 15 minutes on ice. Cell lysates were centrifuged at 20,000xg for 5 minutes. The resulting supernatant (200 µl) was carefully applied onto a 10% to 50% sucrose gradient and centrifuged for 1 hour at 55,000 rpm using a TLS-55 rotor at 4°C with the slowest acceleration and deceleration settings. After the centrifugation, 11 x 200 µl fractions were collected. 40 µl was directly analyzed by immunoblotting as input, while the remainder was diluted to 1 ml with RIPA buffer and immunoprecipitated using rat anti-FLAG beads. The resulting immunoprecipitates were subsequently analyzed by immunoblotting.

To separate the cytosol from ribosomes, as shown in Figures 3B, HEK293T cells (1.5 × 10^6^/well) were seeded onto polylysine-coated 60 cm dishes and transiently transfected with 3µg of FLAG-XBP1u or its mutants and 2µg of HA-Ubiquitin using 20µl of PEI (1mg/ml). After 24 hours of transfection, cells were washed once with 1xPBS and harvested in 1xPBS. The cell pellets were resuspended in 1ml of ribosome buffer and centrifuged for 5 minutes at 20,000xg at 4°C. 850µl of the supernatant was then centrifuged in a 1ml polycarbonate tube at 120,000xg for 40 minutes using a TLA120.2 rotor at 4°C. The supernatant was collected and adjusted to 0.1% SDS and 1% Triton X-100. The ribosomal pellet was washed with 200µl of ice-cold water and resuspended with 650µl of RIPA buffer. Both the cytosol and ribosome fractions were subjected to immunoprecipitation using anti-FLAG beads. The resulting immunoprecipitants were then analyzed by immunoblotting.

#### Sucrose cushion for ribosome isolation

To obtain a ribosome fraction, as depicted in Figure 5B, 6B, and 6C, HEK293T cells or LKO cells (0.8 × 10^6/well) were seeded onto polylysine-coated 6-well plates. Following transfection with siRNA or plasmids, cells were harvested in cold PBS and pelleted by centrifugation at 5,000 g for 5 minutes. The cell pellet was then resuspended in 300 μl of lysis buffer (50 mM Hepes, 500 mM KCl, 10 mM MgCl2, 0.5% Triton, 1 mM DTT, 2.5 U/ml RNasin, 1X Protease Inhibitor). The lysate was left on ice for 15 minutes to completely solubilize proteins, followed by a 20,000 g centrifugation for 10 minutes to clarify. Next, 250 μl of the clarified supernatant was carefully layered onto a 0.5 ml 1M sucrose cushion (consisting of 1M sucrose, 50 mM Hepes, 500 mM KCl, 10 mM MgCl2, and 0.5% Triton) in a 1-ml polycarbonate tube. This was then subjected to ultracentrifugation at 120,000 g for 1 hour using a TLA120.2 rotor. Following centrifugation, the supernatant (containing cytosolic proteins) was gently collected. The ribosome pellet was swiftly rinsed with 200 μl of ice-cold water and resuspended in 100 μl of 1% SDS and 0.1 M Tris-HCl (pH 8.0) at room temperature. The ribosome fraction was then heated for 10 minutes, which was used for further ubiquitination analysis and detecting ribosome-associated nascent chains.

#### In vitro translation and ubiquitination

In vitro transcription and translation were performed using established protocols^24,75^ with the following modifications. PCR products encoding N-terminally FLAG-tagged XBP1u or its mutants were used as templates for in vitro transcription reactions. These PCR products were amplified from pCDNA5/FRT/TO FLAG-XBP1u constructs using a forward primer that included the Sp6 promoter sequence and annealed to the CMV promoter. The reverse primer annealed to the 3’ end of XBP1u, excluding a stop codon for synthesizing XBP1u RNCs. For generating full-length PCR products, a reverse primer annealing to the poly A tail was used. In vitro transcription was performed using SP6 polymerase (New England Biolabs) and RNasin (Promega) at 37°C for 1.5 h. Transcripts lacking a stop codon were directly added to in vitro translation reactions, including hemin, nuclease-treated RRL, S35 methionine (PerkinElmer; #NEG009T005MC), and 10 µM His-tagged ubiquitin (Boston Biochem), and were incubated for 60 min at 32°C. Since transcripts lack a stop codon, ribosomes are stalled and subjected to ubiquitination by the RQC factors as shown previously^32^. The samples were kept on ice and an aliquot was withdrawn for direct analysis as input or denatured using 1% SDS (final concentration). The denatured samples were diluted to 1 ml with IP buffer (50mM Tris-HCl, 150mM NaCl, 1% Triton X-100) and subjected to pull down His-Ubiquitin conjugated XBP1u using Talon beads.

To demonstrate LTN1-mediated ubiquitination of XBP1u RNCs in Figure 4A, after translation for 60 minutes, the reactions were incubated with a buffer containing 20 µM MG132, 10 µM His-Ubiquitin, and an energy regenerating system (1 mM ATP, 1 mM GTP, 12 mM creatine phosphate, 20 µg/ml creatine kinase), or concentrated cytosol prepared from WT HeLa or LKO cells, including 20 µM MG132, 10 µM His-Ubiquitin, and energy regenerating system. The reactions were incubated for 60 minutes at 32°C. The samples were terminated by adjusting to 1% SDS and denatured by boiling, followed by dilution with 1 ml of IP buffer and pull down with Talon beads. For Figure 4B, after the 60-minute translation as above, the translation reactions were untreated, treated with 1 mM Puromycin, 1 µM E1 inhibitor (MLN4924)^54^, or both, as indicated in the Figure 4 legends, and incubated for 30 minutes at 32°C. The samples were denatured, diluted, and pulled down with Talon beads as previously described. For Figure 4C, after puromycin treatment, samples were centrifuged at 120,000xg for 40 minutes using a TLA120.1 rotor before denaturation, dilution, and Talon pull down. For Figure 4D and 4E, the samples were withdrawn at different time points during the puromycin treatment, as indicated in the figure, and processed as above. For Figure 4F, the translation reactions were either left untreated, or treated with Puromycin along with 50µM PR-619. Samples were withdrawn at different time points for Talon pull down, as described above. For S4B, transcripts encoding full-length XBP1-AP* were translated for 60 minutes and incubated with either DMSO or 1 µM E1 inhibitor. The samples were aliquoted for the indicated time points and denatured and enriched for His-Ubiquitin conjugated proteins by Talon pull down. The Talon beads were boiled using 50 µl of 2x SDS sample buffer and analyzed along with inputs by SDS-PAGE followed by autoradiography.

#### Neutral PAGE and Phostag-based immunoblotting

HEK293T cells (0.125 × 10^6^/well) were seeded onto polylysine-coated 24 well plates and transiently transfected with 0.4µg of HA-XBP1u or its mutants using 1µl of Lipofectamine-2000. After 24 hours of transfection, cells were directly harvested in 150ul of 2xSDS buffer prepared using RNAse free water and DTT. The samples were boiled for 10min and analyzed NuPAGE 4 to 12% Bis-Tris gel (Invitrogen) according to the manufacture recommendations. IRE1α phosphorylation was detected by previously described method^76^. Briefly, 5% SDS PAGE gel was made containing 25 μM Phos-tag (Wako). SDS-PAGE was run at 100 V for 2 hr and 40 min. The gel was transferred to nitrocellulose (Bio-Rad, Hercules, CA) and followed with western blotting.

#### Immunofluorescence

293T cells were seeded onto ploy-lysine coated coverslip in a 24-well plate. Twenty-four hours post-transfection with HA-XBP1u or FLAG-XBP1u, cells underwent the following procedure. Cells were fixed using 3.7% formaldehyde for 10 minutes and then permeabilized with 0.1% Triton X-100 for 5 min. Blocking was performed with a blocking buffer (1X PBS including 5% FBS) for 60 min. The primary antibody, rabbit anti-HA or rat anti-Flag, was added at 1:200 dilution in blocking buffer and incubated for 60 minutes. Cells were washed five times for 5 minutes with PBS. Secondary antibodies, rabbit anti-Cy2 or rat anti-Cy2 (Jackson Immune Research), were added at 1:200 dilution in blocking buffer and incubated for 60 minutes, followed by 5 times wash with 5 minutes incubation each time. Cells were then incubated with 5 µg/ml Hoechst stain in 1XPBS for 2 min, washed twice with PBS.

The coverslip was mounted using Fluor mount G (Southern Biotech). Cell images were taken on a Leica Confocal microscope (Leica SP6/SP8) equipped with an HC PL APO 63X oil objective lens and controlled by the Leica Application Suite X software. Image capture settings included 1.0x Zoom, 100 Hz, and a resolution of 1024 x 1024 pixels. For data analysis, the nuclear localization ratio of XBP1 was quantified using image J. The total fluorescence of XBP1u for each cell was first quantified, and the nuclear area was circled and quantified. The nuclear localization ratio was calculated by dividing the nuclear area fluorescence value by the total cell value. Only cells with a clear nucleus were included in the calculation.

#### Quantification and statistical analysis

Quantification of autoradiographs was performed using ImageJ gel analysis. Error bars represent the standard error of the mean (SEM) from two independent experiments, as indicated in the figure legends. For data analysis in Figure 6C, the nuclear localization ratio of XBP1 was quantified using ImageJ, as described above. Data was graphed using GraphPad Prism and represented with a standard error of the mean.

## References

1. Wickner, S., Maurizi, M.R., and Gottesman, S. (1999). Posttranslational quality control: folding, refolding, and degrading proteins. Science 286, 1888–1893. 10.1126/science.286.5446.1888.

2. Ross, C.A., and Poirier, M.A. (2004). Protein aggregation and neurodegenerative disease. Nature Medicine 10, S10–S17. 10.1038/nm1066.

3. Ciechanover, A., and Kwon, Y.T. (2015). Degradation of misfolded proteins in neurodegenerative diseases: therapeutic targets and strategies. Exp Mol Med 47, e147. 10.1038/emm.2014.117.

4. Wang, F., Durfee, L.A., and Huibregtse, J.M. (2013). A cotranslational ubiquitination pathway for quality control of misfolded proteins. Mol Cell 50, 368–378. 10.1016/j.molcel.2013.03.009.

5. Duttler, S., Pechmann, S., and Frydman, J. (2013). Principles of cotranslational ubiquitination and quality control at the ribosome. Mol Cell 50, 379–393. 10.1016/j.molcel.2013.03.010.

6. Sato, S., Ward, C.L., and Kopito, R.R. (1998). Cotranslational ubiquitination of cystic fibrosis transmembrane conductance regulator in vitro. J Biol Chem 273, 7189–7192. 10.1074/jbc.273.13.7189.

7. Turner, G.C., and Varshavsky, A. (2000). Detecting and measuring cotranslational protein degradation in vivo. Science 289, 2117–2120. 10.1126/science.289.5487.2117.

8. Vabulas, R.M., and Hartl, F.U. (2005). Protein synthesis upon acute nutrient restriction relies on proteasome function. Science 310, 1960–1963. 10.1126/science.1121925.

9. Yewdell, J.W., and Nicchitta, C.V. (2006). The DRiP hypothesis decennial: support, controversy, refinement and extension. Trends Immunol 27, 368–373. 10.1016/j.it.2006.06.008.

10. Pechmann, S., Willmund, F., and Frydman, J. (2013). The ribosome as a hub for protein quality control. Mol Cell 49, 411–421. 10.1016/j.molcel.2013.01.020.

11. Princiotta, M.F., Finzi, D., Qian, S.B., Gibbs, J., Schuchmann, S., Buttgereit, F., Bennink, J.R., and Yewdell, J.W. (2003). Quantitating protein synthesis, degradation, and endogenous antigen processing. Immunity 18, 343–354. 10.1016/s1074-7613(03)00051-7.

12. Bengtson, M.H., and Joazeiro, C.A. (2010). Role of a ribosome-associated E3 ubiquitin ligase in protein quality control. Nature 467, 470–473. 10.1038/nature09371.

13. Brandman, O., and Hegde, R.S. (2016). Ribosome-associated protein quality control. Nat Struct Mol Biol 23, 7–15. 10.1038/nsmb.3147.

14. Inada, T. (2020). Quality controls induced by aberrant translation. Nucleic Acids Res 48, 1084–1096. 10.1093/nar/gkz1201.

15. Yip, M.C.J., and Shao, S. (2021). Detecting and Rescuing Stalled Ribosomes. Trends Biochem Sci 46, 731–743. 10.1016/j.tibs.2021.03.008.

16. Brandman, O., Stewart-Ornstein, J., Wong, D., Larson, A., Williams, C.C., Li, G.W., Zhou, S., King, D., Shen, P.S., Weibezahn, J., et al. (2012). A ribosome-bound quality control complex triggers degradation of nascent peptides and signals translation stress. Cell 151, 1042–1054. 10.1016/j.cell.2012.10.044.

17. Schuller, A.P., and Green, R. (2018). Roadblocks and resolutions in eukaryotic translation. Nat Rev Mol Cell Biol 19, 526–541. 10.1038/s41580-018-0011-4.

18. Ito, K., and Chiba, S. (2013). Arrest peptides: cis-acting modulators of translation. Annu Rev Biochem 82, 171–202. 10.1146/annurev-biochem-080211-105026.

19. Wilson, D.N., Arenz, S., and Beckmann, R. (2016). Translation regulation via nascent polypeptide-mediated ribosome stalling. Curr Opin Struct Biol 37, 123–133. 10.1016/j.sbi.2016.01.008.

20. Ingolia, N.T., Lareau, L.F., and Weissman, J.S. (2011). Ribosome profiling of mouse embryonic stem cells reveals the complexity and dynamics of mammalian proteomes. Cell 147, 789–802. 10.1016/j.cell.2011.10.002.

21. Han, P., Shichino, Y., Schneider-Poetsch, T., Mito, M., Hashimoto, S., Udagawa, T., Kohno, K., Yoshida, M., Mishima, Y., Inada, T., and Iwasaki, S. (2020). Genome-wide Survey of Ribosome Collision. Cell Rep 31, 107610. 10.1016/j.celrep.2020.107610.

22. Lu, J., and Deutsch, C. (2008). Electrostatics in the ribosomal tunnel modulate chain elongation rates. J Mol Biol 384, 73–86. 10.1016/j.jmb.2008.08.089.

23. Stein, K.C., Morales-Polanco, F., van der Lienden, J., Rainbolt, T.K., and Frydman, J. (2022). Ageing exacerbates ribosome pausing to disrupt cotranslational proteostasis. Nature 601, 637–642. 10.1038/s41586-021-04295-4.

24. Mariappan, M., Li, X., Stefanovic, S., Sharma, A., Mateja, A., Keenan, R.J., and Hegde, R.S. (2010). A ribosome-associating factor chaperones tail-anchored membrane proteins. Nature 466, 1120–1124. 10.1038/nature09296.

25. Collart, M.A., and Weiss, B. (2020). Ribosome pausing, a dangerous necessity for co-translational events. Nucleic Acids Res 48, 1043–1055. 10.1093/nar/gkz763.

26. Yanagitani, K., Kimata, Y., Kadokura, H., and Kohno, K. (2011). Translational pausing ensures membrane targeting and cytoplasmic splicing of XBP1u mRNA. Science 331, 586–589. 10.1126/science.1197142.

27. Sepulveda, G., Antkowiak, M., Brust-Mascher, I., Mahe, K., Ou, T., Castro, N.M., Christensen, L.N., Cheung, L., Jiang, X., Yoon, D., et al. (2018). Co-translational protein targeting facilitates centrosomal recruitment of PCNT during centrosome maturation in vertebrates. Elife 7. 10.7554/eLife.34959.

28. Zhao, T., Chen, Y.M., Li, Y., Wang, J., Chen, S., Gao, N., and Qian, W. (2021). Disome-seq reveals widespread ribosome collisions that promote cotranslational protein folding. Genome Biol 22, 16. 10.1186/s13059-020-02256-0.

29. Joazeiro, C.A.P. (2019). Mechanisms and functions of ribosome-associated protein quality control. Nat Rev Mol Cell Biol 20, 368–383. 10.1038/s41580-019-0118-2.

30. Howard, C.J., and Frost, A. (2021). Ribosome-associated quality control and CAT tailing. Crit Rev Biochem Mol Biol 56, 603–620. 10.1080/10409238.2021.1938507.

31. D’Orazio, K.N., and Green, R. (2021). Ribosome states signal RNA quality control. Mol Cell 81, 1372–1383. 10.1016/j.molcel.2021.02.022.

32. Shao, S., von der Malsburg, K., and Hegde, R.S. (2013). Listerin-dependent nascent protein ubiquitination relies on ribosome subunit dissociation. Mol Cell 50, 637–648. 10.1016/j.molcel.2013.04.015.

33. Shoemaker, C.J., Eyler, D.E., and Green, R. (2010). Dom34:Hbs1 promotes subunit dissociation and peptidyl-tRNA drop-off to initiate no-go decay. Science 330, 369–372. 10.1126/science.1192430.

34. Pisareva, V.P., Skabkin, M.A., Hellen, C.U., Pestova, T.V., and Pisarev, A.V. (2011). Dissociation by Pelota, Hbs1 and ABCE1 of mammalian vacant 80S ribosomes and stalled elongation complexes. Embo j 30, 1804–1817. 10.1038/emboj.2011.93.

35. Lyumkis, D., Oliveira dos Passos, D., Tahara, E.B., Webb, K., Bennett, E.J., Vinterbo, S., Potter, C.S., Carragher, B., and Joazeiro, C.A. (2014). Structural basis for translational surveillance by the large ribosomal subunit-associated protein quality control complex. Proc Natl Acad Sci U S A 111, 15981–15986. 10.1073/pnas.1413882111.

36. Shen, P.S., Park, J., Qin, Y., Li, X., Parsawar, K., Larson, M.H., Cox, J., Cheng, Y., Lambowitz, A.M., Weissman, J.S., et al. (2015). Protein synthesis. Rqc2p and 60S ribosomal subunits mediate mRNA-independent elongation of nascent chains. Science 347, 75–78. 10.1126/science.1259724.

37. Defenouillère, Q., Yao, Y., Mouaikel, J., Namane, A., Galopier, A., Decourty, L., Doyen, A., Malabat, C., Saveanu, C., Jacquier, A., and Fromont-Racine, M. (2013). Cdc48-associated complex bound to 60S particles is required for the clearance of aberrant translation products. Proc Natl Acad Sci U S A 110, 5046–5051. 10.1073/pnas.1221724110.

38. Shao, S., Brown, A., Santhanam, B., and Hegde, R.S. (2015). Structure and assembly pathway of the ribosome quality control complex. Mol Cell 57, 433–444. 10.1016/j.molcel.2014.12.015.

39. von der Malsburg, K., Shao, S., and Hegde, R.S. (2015). The ribosome quality control pathway can access nascent polypeptides stalled at the Sec61 translocon. Mol Biol Cell 26, 2168–2180. 10.1091/mbc.E15-01-0040.

40. Arakawa, S., Yunoki, K., Izawa, T., Tamura, Y., Nishikawa, S., and Endo, T. (2016). Quality control of nonstop membrane proteins at the ER membrane and in the cytosol. Sci Rep 6, 30795. 10.1038/srep30795.

41. Buchanan, B.W., Mehrtash, A.B., Broshar, C.L., Runnebohm, A.M., Snow, B.J., Scanameo, L.N., Hochstrasser, M., and Rubenstein, E.M. (2019). Endoplasmic reticulum stress differentially inhibits endoplasmic reticulum and inner nuclear membrane protein quality control degradation pathways. J Biol Chem 294, 19814–19830. 10.1074/jbc.RA119.010295.

42. Scavone, F., Gumbin, S.C., Da Rosa, P.A., and Kopito, R.R. (2023). RPL26/uL24 UFMylation is essential for ribosome-associated quality control at the endoplasmic reticulum. Proc Natl Acad Sci U S A 120, e2220340120. 10.1073/pnas.2220340120.

43. Trentini, D.B., Pecoraro, M., Tiwary, S., Cox, J., Mann, M., Hipp, M.S., and Hartl, F.U. (2020). Role for ribosome-associated quality control in sampling proteins for MHC class I-mediated antigen presentation. Proc Natl Acad Sci U S A 117, 4099–4108. 10.1073/pnas.1914401117.

44. Verma, R., Oania, R.S., Kolawa, N.J., and Deshaies, R.J. (2013). Cdc48/p97 promotes degradation of aberrant nascent polypeptides bound to the ribosome. Elife 2, e00308. 10.7554/eLife.00308.

45. Ito-Harashima, S., Kuroha, K., Tatematsu, T., and Inada, T. (2007). Translation of the poly(A) tail plays crucial roles in nonstop mRNA surveillance via translation repression and protein destabilization by proteasome in yeast. Genes Dev 21, 519–524. 10.1101/gad.1490207.

46. Yoshida, H., Matsui, T., Yamamoto, A., Okada, T., and Mori, K. (2001). XBP1 mRNA is induced by ATF6 and spliced by IRE1 in response to ER stress to produce a highly active transcription factor. Cell 107, 881–891. 10.1016/s0092-8674(01)00611-0.

47. Calfon, M., Zeng, H., Urano, F., Till, J.H., Hubbard, S.R., Harding, H.P., Clark, S.G., and Ron, D. (2002). IRE1 couples endoplasmic reticulum load to secretory capacity by processing the XBP-1 mRNA. Nature 415, 92–96. 10.1038/415092a.

48. Yanagitani, K., Imagawa, Y., Iwawaki, T., Hosoda, A., Saito, M., Kimata, Y., and Kohno, K. (2009). Cotranslational targeting of XBP1 protein to the membrane promotes cytoplasmic splicing of its own mRNA. Mol Cell 34, 191–200. 10.1016/j.molcel.2009.02.033.

49. Shanmuganathan, V., Schiller, N., Magoulopoulou, A., Cheng, J., Braunger, K., Cymer, F., Berninghausen, O., Beatrix, B., Kohno, K., von Heijne, G., and Beckmann, R. (2019). Structural and mutational analysis of the ribosome-arresting human XBP1u. Elife 8. 10.7554/eLife.46267.

50. Plumb, R., Zhang, Z.R., Appathurai, S., and Mariappan, M. (2015). A functional link between the co-translational protein translocation pathway and the UPR. Elife 4. 10.7554/eLife.07426.

51. Kanda, S., Yanagitani, K., Yokota, Y., Esaki, Y., and Kohno, K. (2016). Autonomous translational pausing is required for XBP1u mRNA recruitment to the ER via the SRP pathway. Proc Natl Acad Sci U S A 113, E5886–e5895. 10.1073/pnas.1604435113.

52. Chen, C.Y., Malchus, N.S., Hehn, B., Stelzer, W., Avci, D., Langosch, D., and Lemberg, M.K. (2014). Signal peptide peptidase functions in ERAD to cleave the unfolded protein response regulator XBP1u. Embo j 33, 2492–2506. 10.15252/embj.201488208.

53. Azzam, M.E., and Algranati, I.D. (1973). Mechanism of puromycin action: fate of ribosomes after release of nascent protein chains from polysomes. Proc Natl Acad Sci U S A 70, 3866–3869. 10.1073/pnas.70.12.3866.

54. Soucy, T.A., Smith, P.G., Milhollen, M.A., Berger, A.J., Gavin, J.M., Adhikari, S., Brownell, J.E., Burke, K.E., Cardin, D.P., Critchley, S., et al. (2009). An inhibitor of NEDD8-activating enzyme as a new approach to treat cancer. Nature 458, 732–736. 10.1038/nature07884.

55. Altun, M., Kramer, H.B., Willems, L.I., McDermott, J.L., Leach, C.A., Goldenberg, S.J., Kumar, K.G., Konietzny, R., Fischer, R., Kogan, E., et al. (2011). Activity-based chemical proteomics accelerates inhibitor development for deubiquitylating enzymes. Chem Biol 18, 1401–1412. 10.1016/j.chembiol.2011.08.018.

56. Hessa, T., Sharma, A., Mariappan, M., Eshleman, H.D., Gutierrez, E., and Hegde, R.S. (2011). Protein targeting and degradation are coupled for elimination of mislocalized proteins. Nature 475, 394–397. 10.1038/nature10181.

57. Hu, X., Wang, L., Wang, Y., Ji, J., Li, J., Wang, Z., Li, C., Zhang, Y., and Zhang, Z.R. (2020). RNF126-Mediated Reubiquitination Is Required for Proteasomal Degradation of p97-Extracted Membrane Proteins. Mol Cell 79, 320–331.e329. 10.1016/j.molcel.2020.06.023.

58. Rodrigo-Brenni, M.C., Gutierrez, E., and Hegde, R.S. (2014). Cytosolic quality control of mislocalized proteins requires RNF126 recruitment to Bag6. Mol Cell 55, 227–237. 10.1016/j.molcel.2014.05.025.

59. Banaszynski, L.A., Chen, L.C., Maynard-Smith, L.A., Ooi, A.G., and Wandless, T.J. (2006). A rapid, reversible, and tunable method to regulate protein function in living cells using synthetic small molecules. Cell 126, 995–1004. 10.1016/j.cell.2006.07.025.

60. Yoshida, H., Uemura, A., and Mori, K. (2009). pXBP1(U), a negative regulator of the unfolded protein response activator pXBP1(S), targets ATF6 but not ATF4 in proteasome-mediated degradation. Cell Struct Funct 34, 1–10. 10.1247/csf.06028.

61. Li, X., Sun, S., Appathurai, S., Sundaram, A., Plumb, R., and Mariappan, M. (2020). A Molecular Mechanism for Turning Off IRE1α Signaling during Endoplasmic Reticulum Stress. Cell Rep 33, 108563. 10.1016/j.celrep.2020.108563.

62. Thrower, J.S., Hoffman, L., Rechsteiner, M., and Pickart, C.M. (2000). Recognition of the polyubiquitin proteolytic signal. Embo j 19, 94–102. 10.1093/emboj/19.1.94.

63. Thrun, A., Garzia, A., Kigoshi-Tansho, Y., Patil, P.R., Umbaugh, C.S., Dallinger, T., Liu, J., Kreger, S., Patrizi, A., Cox, G.A., et al. (2021). Convergence of mammalian RQC and C-end rule proteolytic pathways via alanine tailing. Mol Cell 81, 2112–2122.e2117. 10.1016/j.molcel.2021.03.004.

64. Tesina, P., Ebine, S., Buschauer, R., Thoms, M., Matsuo, Y., Inada, T., and Beckmann, R. (2023). Molecular basis of eIF5A-dependent CAT tailing in eukaryotic ribosome-associated quality control. Mol Cell 83, 607–621.e604. 10.1016/j.molcel.2023.01.020.

65. Sitron, C.S., and Brandman, O. (2019). CAT tails drive degradation of stalled polypeptides on and off the ribosome. Nat Struct Mol Biol 26, 450–459. 10.1038/s41594-019-0230-1.

66. Zhang, Z.R., Bonifacino, J.S., and Hegde, R.S. (2013). Deubiquitinases sharpen substrate discrimination during membrane protein degradation from the ER. Cell 154, 609–622. 10.1016/j.cell.2013.06.038.

67. Helenius, A., and Aebi, M. (2004). Roles of N-linked glycans in the endoplasmic reticulum. Annu Rev Biochem 73, 1019–1049. 10.1146/annurev.biochem.73.011303.073752.

68. Lu, Y., Lee, B.H., King, R.W., Finley, D., and Kirschner, M.W. (2015). Substrate degradation by the proteasome: a single-molecule kinetic analysis. Science 348, 1250834. 10.1126/science.1250834.

69. Pierce, N.W., Kleiger, G., Shan, S.O., and Deshaies, R.J. (2009). Detection of sequential polyubiquitylation on a millisecond timescale. Nature 462, 615–619. 10.1038/nature08595.

70. Eshraghi, M., Karunadharma, P.P., Blin, J., Shahani, N., Ricci, E.P., Michel, A., Urban, N.T., Galli, N., Sharma, M., Ramírez-Jarquín, U.N., et al. (2021). Mutant Huntingtin stalls ribosomes and represses protein synthesis in a cellular model of Huntington disease. Nat Commun 12, 1461. 10.1038/s41467-021-21637-y.

71. Yang, J., Hao, X., Cao, X., Liu, B., and Nyström, T. (2016). Spatial sequestration and detoxification of Huntingtin by the ribosome quality control complex. Elife 5. 10.7554/eLife.11792.

72. Mali, P., Yang, L., Esvelt, K.M., Aach, J., Guell, M., DiCarlo, J.E., Norville, J.E., and Church, G.M. (2013). RNA-guided human genome engineering via Cas9. Science 339, 823–826. 10.1126/science.1232033.

73. Ran, F.A., Hsu, P.D., Wright, J., Agarwala, V., Scott, D.A., and Zhang, F. (2013). Genome engineering using the CRISPR-Cas9 system. Nat Protoc 8, 2281–2308. 10.1038/nprot.2013.143.

74. Labun, K., Montague, T.G., Krause, M., Torres Cleuren, Y.N., Tjeldnes, H., and Valen, E. (2019). CHOPCHOP v3: expanding the CRISPR web toolbox beyond genome editing. Nucleic Acids Res 47, W171–w174. 10.1093/nar/gkz365.

75. Sharma, A., Mariappan, M., Appathurai, S., and Hegde, R.S. (2010). In vitro dissection of protein translocation into the mammalian endoplasmic reticulum. Methods Mol Biol 619, 339–363. 10.1007/978-1-60327-412-8_20.

76. Yang, L., Xue, Z., He, Y., Sun, S., Chen, H., and Qi, L. (2010). A Phos-tag-based approach reveals the extent of physiological endoplasmic reticulum stress. PLoS One 5, e11621. 10.1371/journal.pone.0011621.

