## Supplemental Figures for "Nascent Chain Ubiquitination is Uncoupled from Degradation to Enable Protein Maturation"

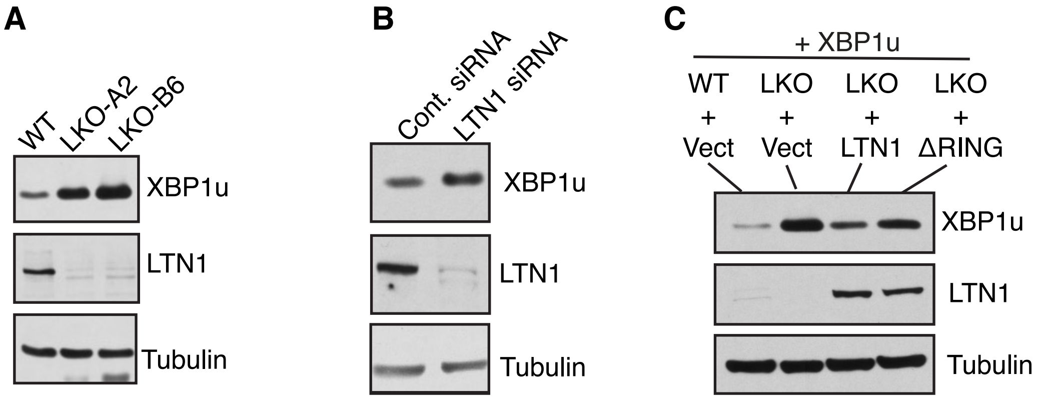


**Figure S1: XBP1u accumulates in cells depleted of LTN1 E3 ligase activity**
(A) Wild type (WT) 293T or different clones of LKO cells were transfected with XBP1u and analyzed by immunoblotting.
(B) Control siRNA or LTN1 siRNA treated cells were transfected with XBP1u and analyzed by immuno-blotting.
(C) Cells expressing the indicated constructs were analyzed by immunoblotting.

Related to **Figure 1**

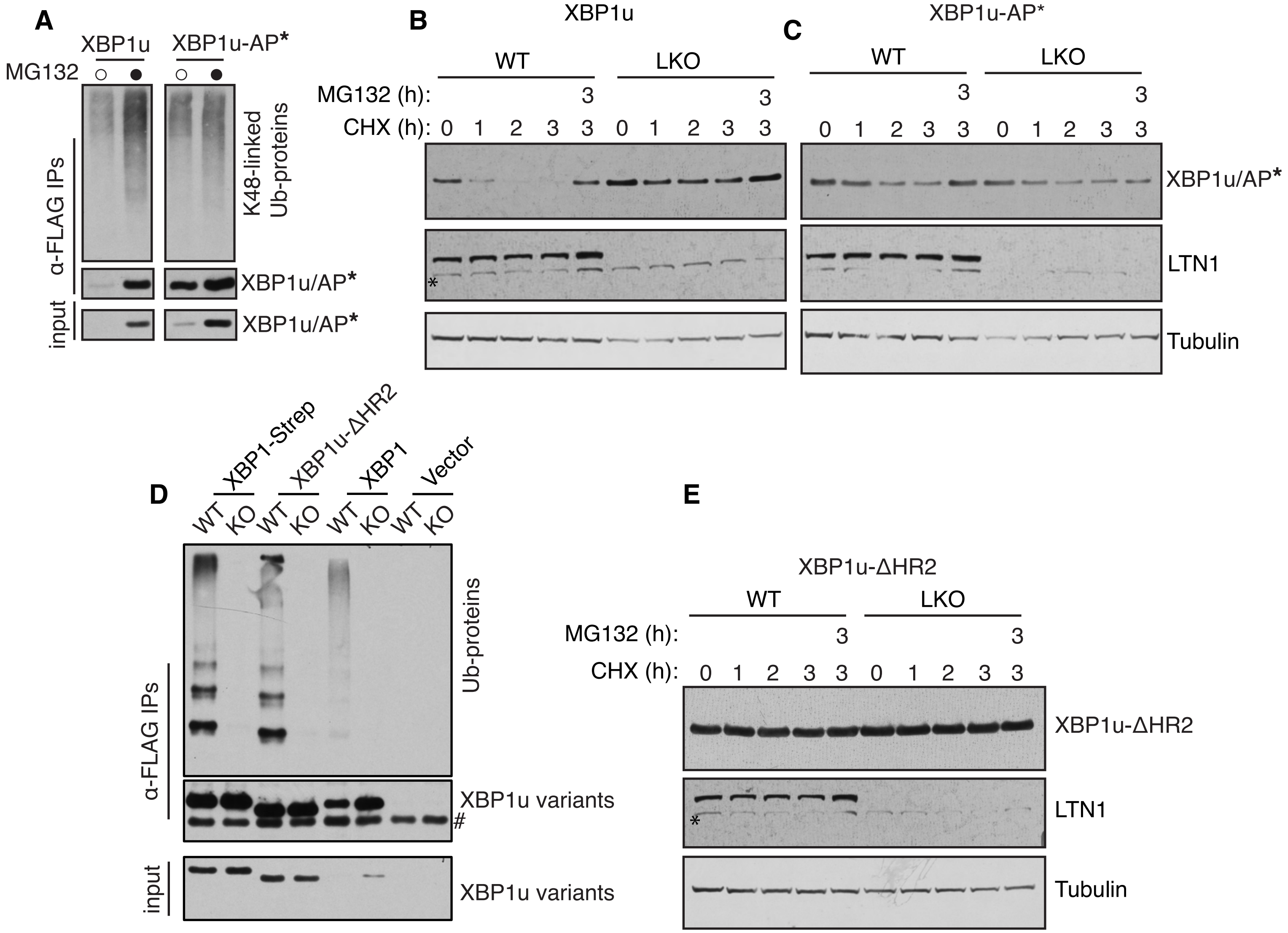


**Figure S2: Nascent chain ubiquitination by LTN1 on stalled ribosomes does not mediate proteasomal degradation.**
(A) FLAG-XBP1u variants were transfected along with HA-ubiquitin into 293T. The resulting cell lysates were immunoprecipitated with anti-FLAG beads and probed for XBP1u with an anti-FLAG and an K48-linked ubiquitin chain specific antibody.
(B) XBP1u-expressing wild-type (WT) cells or LTN1 knockout (LKO) cells were treated with cycloheximide (100 $\mu$g/ml) alone or with 20 $\mu$M MG132. Cells were then harvested at the specified time points and subjected to immunoblotting analysis. * unspecific band.
(C) XBP1u-AP* expressing cells were treated and analyzed as in (B).
(D) Empty vector or FLAG-tagged XBP1u variants were transfected along with HA-ubiquitin into WT or LKO cells, followed by anti-FLAG immunoprecipitations (IPs). The resulting samples were probed for ubiquitinated proteins (Ub-proteins) using an HA antibody, and for XBP1u variants with an anti-FLAG antibody. # Background band.
(E) XBP1u-ΔHR2 expressing WT and LKO cells were treated and analyzed as in (B). * unspecific band.

Related to **Figure 2**

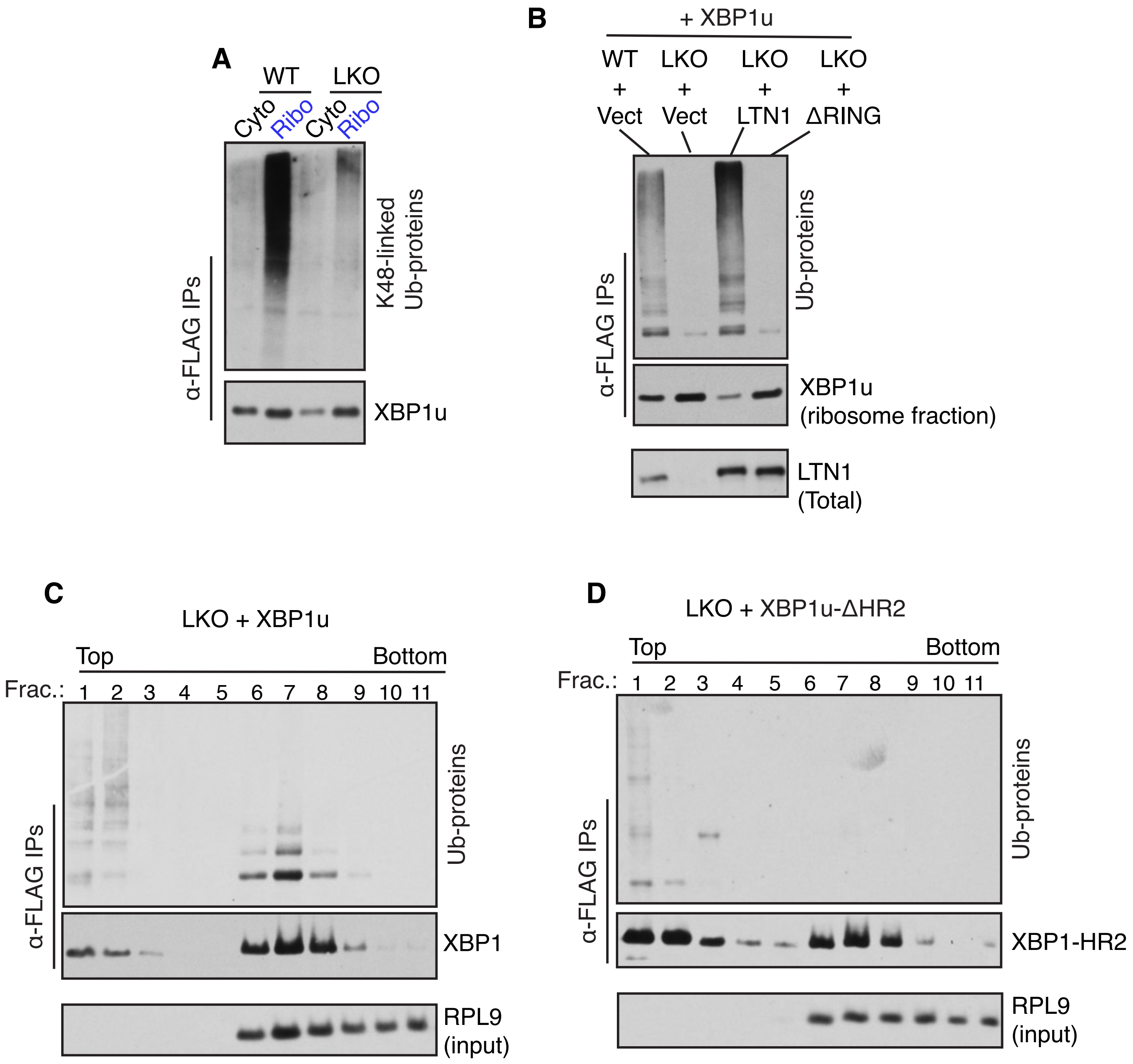


**Figure S3: Ubiquitination of XBP1u on stalled ribosomes requires LTN1 E3 ligase activity**
(A) WT and LKO cells expressing XBP1u were separated into cytosol and ribosome fractions. The resulting samples were immunoprecipitated using anti-FLAG beads and probed for XBP1u conjugated with K48-linked ubiquitin chains.
(B) Ribosomes were isolated from the indicated cells co-expressing FLAG-XBP1u and HA-ubiquitin along with untagged LTN1 or LTN1ΔRING. The ribosome pellets were denatured, diluted, and immunoprecipitated for XBP1u using anti-FLAG antibody. The resulting immunoprecipitates were immunoblotted for XBP1u with an anti-FLAG antibody and ubiquitinated proteins using an HA antibody.
(C, D) LKO cell lysates of XBP1u or XBP1uΔHR2 along with HA-ubiquitin were subjected to a 10-50% sucrose density gradient. Input fractions were directly probed for the ribosomal protein RPL9, while XBP1u and its mutant were immunoprecipitated using anti-FLAG beads and analyzed by immunoblotting.

Related to **Figure 3**

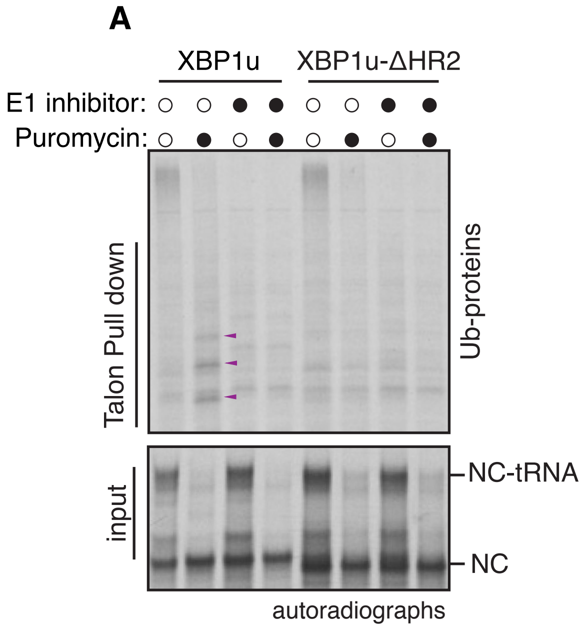


**Figure S4: Continuous ubiquitination of nascent chain is necessary to compete against cytosolic deubiquitinates (DUBs).**

(A) Ubiquitin modified RNCs of XBP1u and its mutant were prepared in the rabbit reticulocyte lysate (RRL) based in vitro translation system including Hi-Ubiquitin and 35S-methionine. Subsequently, the reactions were treated with puromycin for 45 minutes. An E1 inhibitor (MLN4924) was included without or with puromycin. The samples were then denatured and either directly analyzed as input or subjected to Talon pull-down to capture His-Ubiquitin-conjugated radiolabeled XBP1u. The resulting samples were visualized by autoradiography. NC: nascent chain. Arrow head: re-ubiquitination.

(B) Transcripts encoding full length XBP1u-AP* translated in RRL for 45min and treated with either DMSO or E1 inhibitor (MLN4924) and then sampled at the specified time intervals. The samples were denatured and analyzed as described in (A).

Related to **Figure 4**

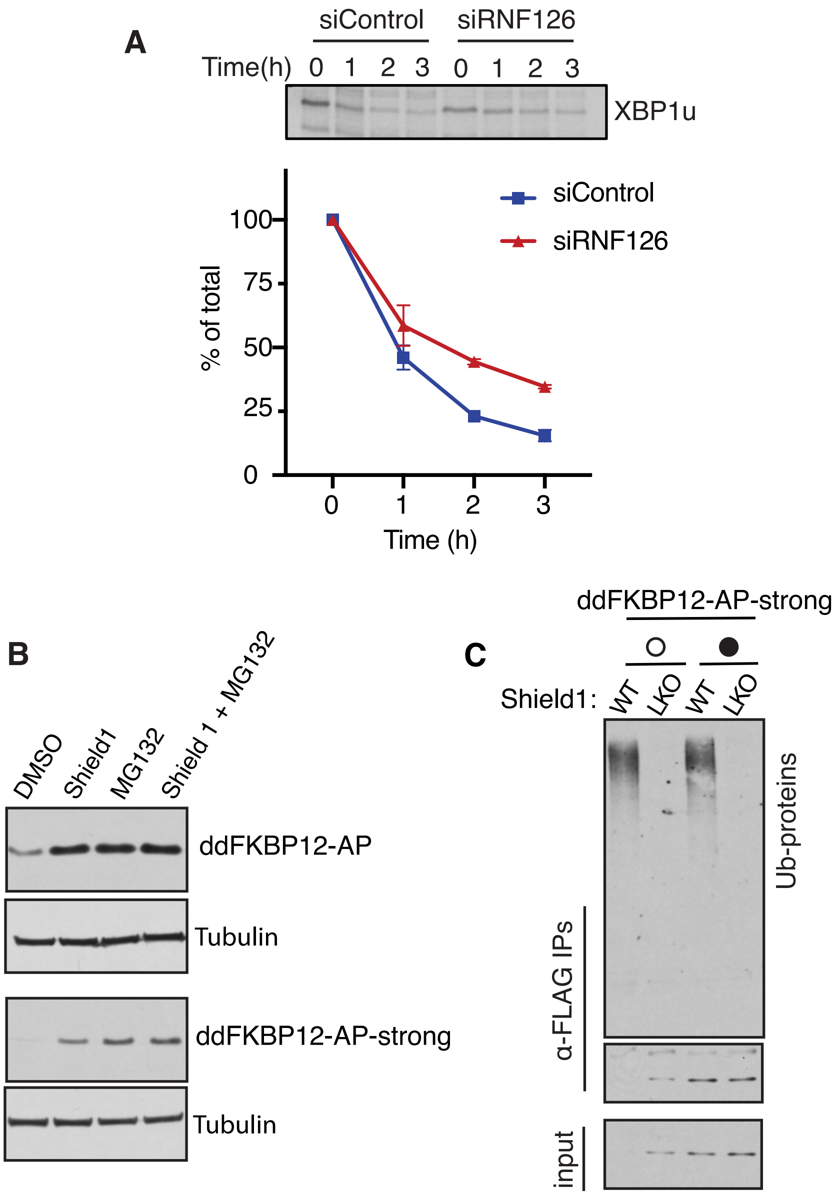


**Figure S5: Re-ubiquitination of nascent chains exposing a degron depends on cytosolic quality control.**

(A) Pulse-chase analysis of Control or RNF126 siRNA treated cells expressing XBP1u. The graph represents data from two independent experiments and is presented as means ± SEMs.

(B) Cells expressing either dd FKBP12-AP or dd FKBP12-AP-strong, generated introducing S255A mutation in the arrest peptide (AP), were treated with 1 $\mu$M Shield1, 20 $\mu$M MG132, or both and analyzed by immunoblotting.

(C) WT or LKO cells expressing HA-ubiquitin and dd FKBP12-AP-strong were untreated or treated with Shield1. The cell lysates were immunoprecipitated and analyzed by immunoblotting.

Related to **Figure 5**

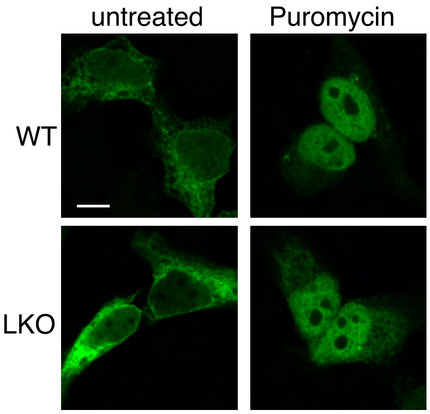


**Figure S6: Puromycin treatment releases XBP1u nascent chains from stalled ribosomes in the ER.**
WT or LKO cells expressing XBP1u were treated with 250 $\mu$M puromycin for 30 min. Subsequently, cells were processed using an immunostaining procedure to label XBP1u (green) with rat anti-FLAG. Scale bar is 10 $\mu$m.

Related to **Figure 6**
